# Phosphatidylinositol-4-phosphate signaling regulates dense granule biogenesis and exocytosis in *Toxoplasma gondii*

**DOI:** 10.1101/2023.01.09.523261

**Authors:** Angela Arabiotorre, Megan Formanowicz, Vytas A. Bankaitis, Aby Grabon

## Abstract

Phosphoinositide metabolism defines the foundation of a major signaling pathway that is conserved throughout the eukaryotic kingdom. The 4-OH phosphorylated phosphoinositides such as phosphatidylinositol-4-phosphate (PtdIns4P) and phosphatidylinositol-4,5-bisphosphate are particularly important molecules as these execute intrinsically essential activities required for the viability of all eukaryotic cells studied thus far. Using intracellular tachyzoites of the apicomplexan parasite *Toxoplasma gondii* as model for assessing primordial roles for PtdIns4P signaling, we demonstrate the presence of PtdIns4P pools in Golgi/trans-Golgi (TGN) system and in post-TGN compartments of the parasite. Moreover, we show that deficits in PtdIns4P signaling result in structural perturbation of compartments that house dense granule cargo with accompanying deficits in dense granule exocytosis. Taken together, the data report a direct role for PtdIns4P in dense granule biogenesis and exocytosis. The data further indicate that the biogenic pathway for secretion-competent dense granule formation in *T. gondii* is more complex than simple budding of fully matured dense granules from the TGN.

## INTRODUCTION

Phosphoinositides (PIPs) are phosphorylated derivatives of the glycerol-based phospholipid phosphatidylinositol (PtdIns). PIPs are chemically distinguished by positionally specific phosphorylations at the 3-OH, 4-OH and 5-OH positions on the inositol ring that constitutes the headgroup of these lipids (Carpenter and Cantley, 1990). The relatively low chemical complexity of the PIP cohort notwithstanding (mammals express seven PIPs while yeast produce four) these molecules regulate literally hundreds of distinct biological outcomes (reviewed in Di Paolo and Camilli, 2006; Balla, 2013). Among the various cellular events regulated PIPs, maintenance of organelle identity and regulation of membrane trafficking events is of direct relevance to this study. In yeast and higher eukaryotes, PIP-regulated membrane trafficking processes include: (1) endocytosis and the formation of PtdIns(3)P-positive vesicles from the PM and its fusion with the endocytic system (Gillooly et al., 2000; Petiot et al., 2003; Ikonomov et al., 2006: Whitley et al., 2009; Duex et al., 2006), (2) the PtdIns(3)P-dependent formation of autophagosomes and PtdIns(3,5)P_2_-dependent fusion with lysosomes (Mizushima et al., 2008), and (3) formation of exocytic vesicles at the TGN in a PtdIns4P-dependent manner and their subsequent transport to the plasma membrane (Rivas et al., 1999; Hama et al., 1999; Walch-Solimena et al., 1999 Jović et al., 2012). Considerable effort has been productively invested in the identification and characterization of protein factors that function as downstream effectors of PIP signaling in these various membrane trafficking events – primarily in yeast and in mammalian cells (Rothman et al., 1994; Schekman et al., 1996; Olayioye et al., 2019). Very little is known about the relationship of PIP signaling and control of membrane trafficking in ancient eukaryotic systems, however.

Organisms of the phylum Apicomplexa are primitive eukaryotes that exhibit an obligate intracellular parasitic lifestyle that comes with unique adaptations. Given the deep roots of apicomplexan parasites in the eukaryotic evolutionary tree (Escalante et al., 1995; Levine et al., 2018), these parasites offer exceptional models for understanding the primordial design of PIP signaling systems. *T. gondii* is arguably the most experimentally tractable apicomplexan parasite. This organism can be cultured in vitro, is amenable to genetic manipulation, and rodent models have been established for studying *T. gondii* infection (Pittman et al., 2014; Jacot and Soldati-Favre, 2020). Like other apicomplexans, *T. gondii* harbors unique endomembrane organelles that are dedicated to the execution of activities essential for completion of their complex parasitic life cycles. For instance, a set of specialized late secretory organelles – micronemes, rhoptries, and dense granules (DGs) – are used to store and exocytose cargos required for successful invasion, intracellular survival, and egress from host cells (Carruthers et al., 1999; Nichols et al., 1983; Bai et al., 2018). Although some factors involved in the biogenesis and function of micronemes and rhoptries have been characterized, very little is known concerning mechanisms of DG biogenesis. Moreover, the roles of PIPs in the formation of, and cargo trafficking from, these organelles remain unstudied.

PIP binding domains have been successfully used to localize and analyze intracellular PIP pools applied in many cell types ranging from yeast to plants and to mammals. Such approaches have also been applied to *T. gondii* to visualize intracellular distributions of PtdIns3P and PtdIns(3,5)P_2_ in this parasite. Those studies revealed important roles for the 3-OH PIPs in the biogenesis and/or inheritance of the apicoplast (Tawk et al., 2011; Daher et al., 2015). However, the biological function of the 4’-OH phosphorylated PIPs, particularly PdIns4P, has not been studied in detail in neither *T. gondii* nor in Apicomplexa in general – although a Golgi pool of PtdIns4P has been reported in *Plasmodium falciparum* (McNamara et al., 2013).

Herein, we use the PtdIns4P-binding domains of mammalian GOLPH3, FAPP1 and bacterial SidM (P4M) proteins to obtain spatial and functional information regarding PtdIns4P signaling in intracellular tachyzoites of *T. gondii*. Our data demonstrate the presence of PtdIns4P pools in the Golgi/TGN system and post-TGN compartments and provide evidence for cell-cycle-dependent regulation of PtdIns4P signaling. Moreover, we show that compromised PtdIns4P signaling results in structural perturbation of DGs with accompanying deficits in exocytosis of DG cargo. Taken together, the data not only demonstrate that PtdIns4P itself is a key regulator of DG biogenesis and exocytosis, but also suggest that the biogenic pathway for DG formation in *T. gondii* is more complex than simple budding from the TGN -- as is presently proposed (Griffith et al., 2022). Finally, the collective results argue for a primordial role for PtdIns4P signaling in membrane trafficking through the late stages of the secretory pathway – a role that has been preserved throughout the *Eukaryota* from yeast to primates.

## RESULTS

### Visualization of *T. gondii* PtdIns4P pools by genetically encoded biosensors

To visualize pools of PtdIns4P and its higher-order derivatives PtdIns(4,5)P_2_ and PtdIns(3,4)P_2_ in *T. gondii*, fluorescent biosensors based on PIP binding domains of known specificity were expressed in the parasite and imaged. A tandem human PLCδ PH domain fused to enhanced green fluorescent protein at the C-terminus (2xPHPLCδ-EGFP; (Várnai et al., 1998), and a PH domain of human TAPP1 fused to YFP at the N-terminus (YFP-TAPP1PH; Hogan et al., 2004), were used to monitor distribution of PtdIns(4,5)P_2_ and its regioisomer PtdIns(3,4)P_2_, respectively. Expression of these biosensors was driven by the tubulin (Tub1) promoter. The corresponding reporter genes were transfected into *T. gondii* RH tachyzoites and their profiles recorded under a transient transfection regime. Wide-field fluorescence imaging performed 24 hours post-transfection (hpt) reported recruitment of both biosensors to the plasma membrane of intracellular parasites although diffuse cytoplasmic signals were also detected in both cases (**Figure 1A**). These data suggest both PIP_2_ species localize predominantly to the plasma membrane of *T. gondii*. These profiles are consistent with those reported in other eukaryotes (Hammond et al., 2012; Halet et al., 2002; Heo et al., 2006).

**Figure 1.**
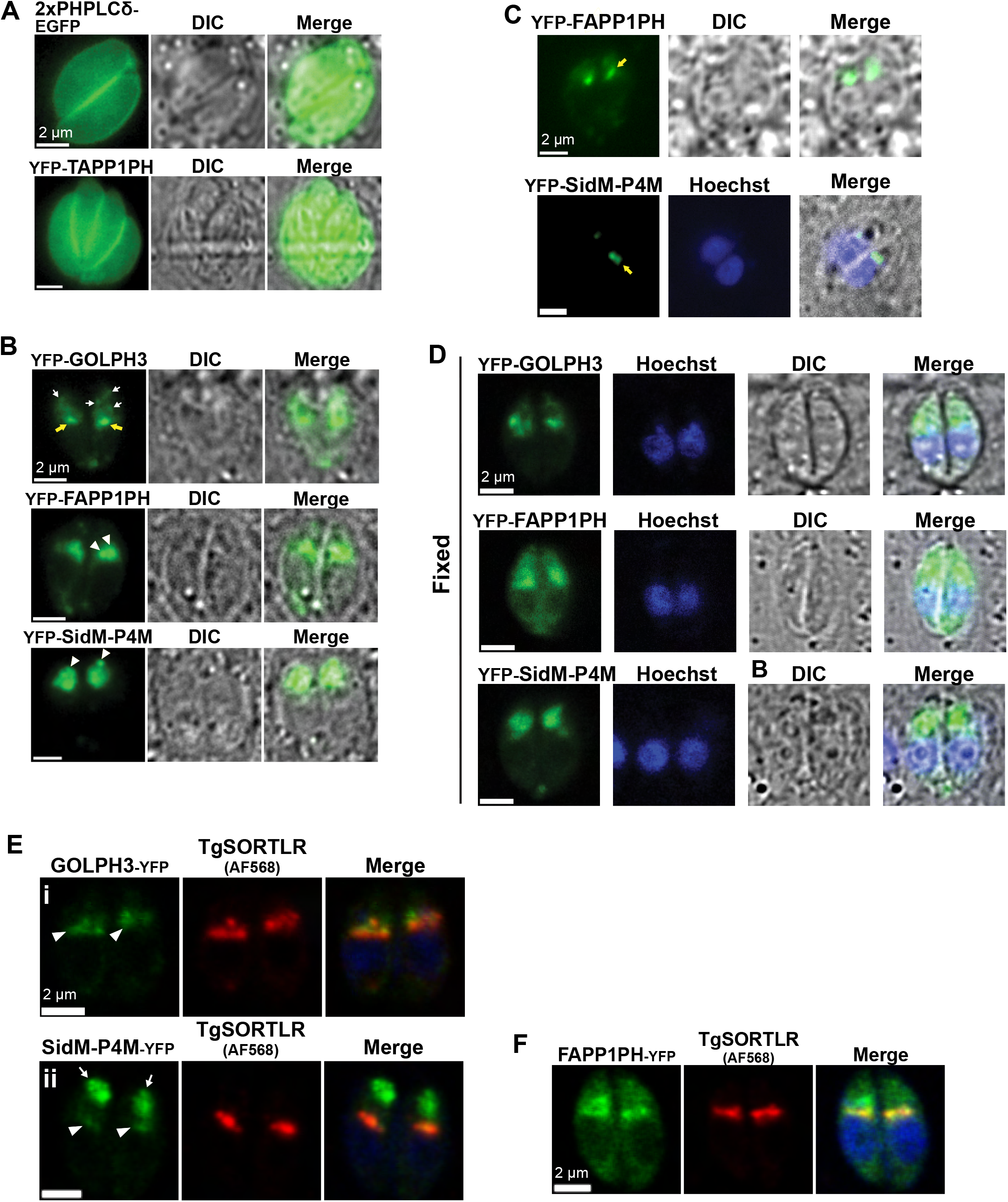
Imaging of PtdIns(4,5)P_2_ and PtdIns(3,4)P_2_ at the plasma membrane and PtdIns4P pools at Golgi/TGN and post-TGN compartments using specific biosensors. **(A-B)** Widefield fluorescence microscopy of live intracellular parasites expressing the indicated PIP biosensors are shown. **(A)** PtdIns(4,5)P_2_ (2xPHPLCδ-EGFP) and PtdIns(3,4)P_2_ (YFP-TAPP1PH) localize to the plasma membrane. **(B)** PtdIns4P pools in RH-GOLPH3 cells localize to a single stack-like structure (yellow arrows) and scattered anterior small vesicles (white arrows); PtdIns4P pools in RH-FAPP1PH and RH-SidM-P4M strains localize to the LAP body (arrowheads). **(C)** RH-FAPP1PH and RH-SidM-P4M cells expressing low levels of the respective biosensors show PtdIns4P in a single stack-like structure (yellow arrows). **(D)** Fluorescence images of biosensor localization in RH-GOLPH3, RH-FAPP1PH and RH-SidM-P4M strains. In RH-GOLPH3 and RH-SidM-P4M parasites. The localization of the biosensor is conserved following fixation, while RH-FAPP1PH cells show a more diffused pattern when compared to the profiles seen in live cells. Parasite DNA is visualized using DAPI (blue). **(E-F)** Confocal images of immunofluorescence analysis (IFA) for the identification of Golgi/TGN (using anti-TgSORTLR immunoglobulin) in parasites during G1/S phase. **(E)** GOLPH3-YFP and SidM-P4M-YFP colocalize with TgSORTLR (arrowheads). The LAP body forms in a post-Golgi/TGN compartment in RH-SidM-P4M cells (arrows), but not in RH-GOLPH3 cells. **(F)** The LAP body forms in a post-Golgi/TGN compartment in RH-FAPP1PH cells.

PtdIns4P is a metabolic precursor for both PtdIns(4,5)P_2_ and PtdIns(3,4)P_2_ and serves intrinsically essential signaling functions in all eukaryotic cells. To monitor PtdIns4P distribution in *T. gondii*, three independent biosensors were designed that exploit distinct PtdIns4P-specific binding modules. These included: (i) the N-terminus of human GOLPH3, (ii) the PH domain of human FAPP1 that we refer to as FAPP1PH, and (iii) the P4M domain of the *Legionella pneumophilia* SidM protein (Wood et al., 2009; Dowler et al., 2000, Del Campo et al., 2014).

These reporters were chosen because of their abilities to detect distinct, but overlapping, pools of PtdIns4P in secretory pathway organelles of mammalian cells and yeast (Levine and Munro, 2002; Orii et al., 2021; Xie et al., 2018; Hammond et al., 2014). Moreover, although these domains are all specific for PtdIns4P binding, these differ in their affinities for PtdIns4P. For example, the GOLPH3 module exhibits a relatively modest PtdIns4P-binding affinity (K_D_ = 2.6 ± 0.2 μM) (Wood et al., 2009), whereas the SidM P4M domain has the highest (K_D_ = 3.8 ± 2.7 nM) (Del Campo et al., 2014). The human FAPP1PH domain exhibits an intermediate binding affinity that is sensitive to pH (K_D_ = 200 nM at pH 6.5 and 460 nM at pH 7.4; He et al., 2011). The transfection and imaging regime for these experiments was as described above for the PtdIns(4,5)P_2_ and PtdIns(3,4)P_2_ biosensors.

In describing the results, all transiently transfected RH-WT parasites are heretofore referred to by the convention of RH followed by the designation of the corresponding PtdIns4P-binding domain -- i.e., RH-GOLPH3, RH-FAPP1PH and RH-SidM-P4M. Unless otherwise stated, all cells analyzed had not yet initiated the process of cell division by endodyogeny and are broadly described as G1/S cells. Wide-field fluorescence imaging reported multiple PtdIns4P distribution profiles in G1/S cells as a function of the specific biosensor used to image PtdIns4P profiles (**Figure 1B**). For example, the RH-GOLPH3 biosensor highlighted a single juxta-nuclear stack and punctate structures that showed a predominantly apical distribution.

These apical structures were visualized as a single large and intensely stained puncta in both RH-FAPP1PH and RH-SidM-P4M parasites -- especially in RH-SidM-P4M parasites (**Figure 1B**, arrowheads). For the remainder of this work, we refer to these apical structures as LAP bodies (<u>L</u>arge <u>A</u>nterior <u>P</u>unctate body). That the appearance of LAP bodies reflected the affinity of the PtdIns4P biosensor was indicated by the following observations. First, expression of the higher affinity FAPP1PH- and SidM-P4M-based biosensors resulted in appearance of LAP bodies, whereas expression of the lower affinity GOLPH3-based biosensor did not. Second, parasites expressing lower fluorescence intensities for the FAPP1PH and SidM-P4M reporters showed distributions that recapitulated those of GOLPH3 biosensor-expressing parasites (**Figure 1C**). We also noted that all three PtdIns4P biosensors marked small cytoplasmic puncta that were frequently in the vicinity of the residual body in tachyzoites that had completed the first round of cell division post-infection.

### Golgi/TGN and post-TGN compartments host PtdIns4P pools

The identities of PtdIns4P-containing organelles were determined in double-label immunofluorescence analyses using antibodies directed against established markers for specific parasite compartments. In these efforts, we first assessed whether the localization of PtdIns4P biosensors in parasites fixed with 4% paraformaldehyde (PFA) recapitulated the profiles observed in live cells. Indeed, the localization profiles of GOLPH3 and SidM-P4M domains in fixed parasites were consistent with those observed in live cells (**Figure 1D**).

However, while the juxta-nuclear compartment localization and LAP body were both detected in fixed RH-FAPP1PH parasites, an unexpected diffuse cytosolic signal was also seen (**Figure 1D**). This fixation artifact had been previously noted in mammalian cells for the FAPP1PH domain (Schmiedeberg et al., 2009). Therefore, further use of the RH-FAPP1PH was limited to live parasite experiments whereas the RH-SidM-P4M reporter was deployed in immunofluorescence studies using fixed parasites.

As significant pools of PtdIns4P are present in Golgi/trans-Golgi network (TGN) membranes in mammalian and yeast cells (Clayton et al., 2013; Rivas et al., 1999; Hama et al., 1999), immunofluorescence approaches were used to assess RH-GOLPH3 and RH-SidM-P4M co-localization with the *T. gondii* TGN marker TgSORTLR (Sortilin-like Receptor; Sloves et al., 2012). Both PtdIns4P biosensors colocalized with varying degrees with TgSORTLR. The GOLPH3-derived PtdIns4P sensor localized to the Golgi/TGN membranes (Pearson’s correlation coefficient ± SEM = 0.62 ± 0.03; **Figure 1Ei**, arrowheads). Recruitment of SidM-P4M to the Golgi/TGN compartment was less efficient but nonetheless detectable (Pearson’s correlation coefficient ± SEM = 0.14 ± 0.04; **Figure 1Eii**, arrowheads). This relatively low incidence of colocalization was likely due to SidM-P4M being strongly recruited to a distinct PtdIns4P pool that persists in a post-TGN compartment intracellular compartment – i.e. the LAP body (**Figure 1Eii**, arrows).

Although colocalization of FAPP1PH with Golgi/TGN membranes was not quantified due to the fixation artifacts described above, this biosensor highlighted both Golgi/TGN membranes and the LAP body in a manner similar to that observed for the SidM-P4M biosensor (**Figure 1F**). Taken together, these data confirm that the stacked juxta-nuclear compartment to which the PtdIns4P biosensors localized represented the Golgi/TGN system. The data further identified a distinct PtdIns4P pool that resided in a second intracellular compartment (the LAP body) characterized by its large punctate profile and its localization to the apical region of the cell.

### The LAP body is a post-Golgi compartment that harbors GRA3 secretory cargo

What is the nature of the LAP body that houses the second PtdIns4P pool identified by the SidM-P4M biosensor? The most attractive possibility was that this structure represented a secretory organelle. To examine this possibility in further detail, we assessed co-localization of RH-SidM-P4M with the microneme marker MIC3 and with the genetically-encoded rhoptry marker ROP1-RFP (**Figure 2A**). As illustrated by the images in **Figure 2B**, the SidM-P4M biosensor failed to colocalize with either the microneme or the rhoptry marker. Moreover, the morphologies of these secretory organelles were not obviously disturbed in SidM-P4M-expressing parasites when compared to RH-WT stained for MIC3 and to RH-ROP1-RFP parasites, respectively.

**Figure 2.**
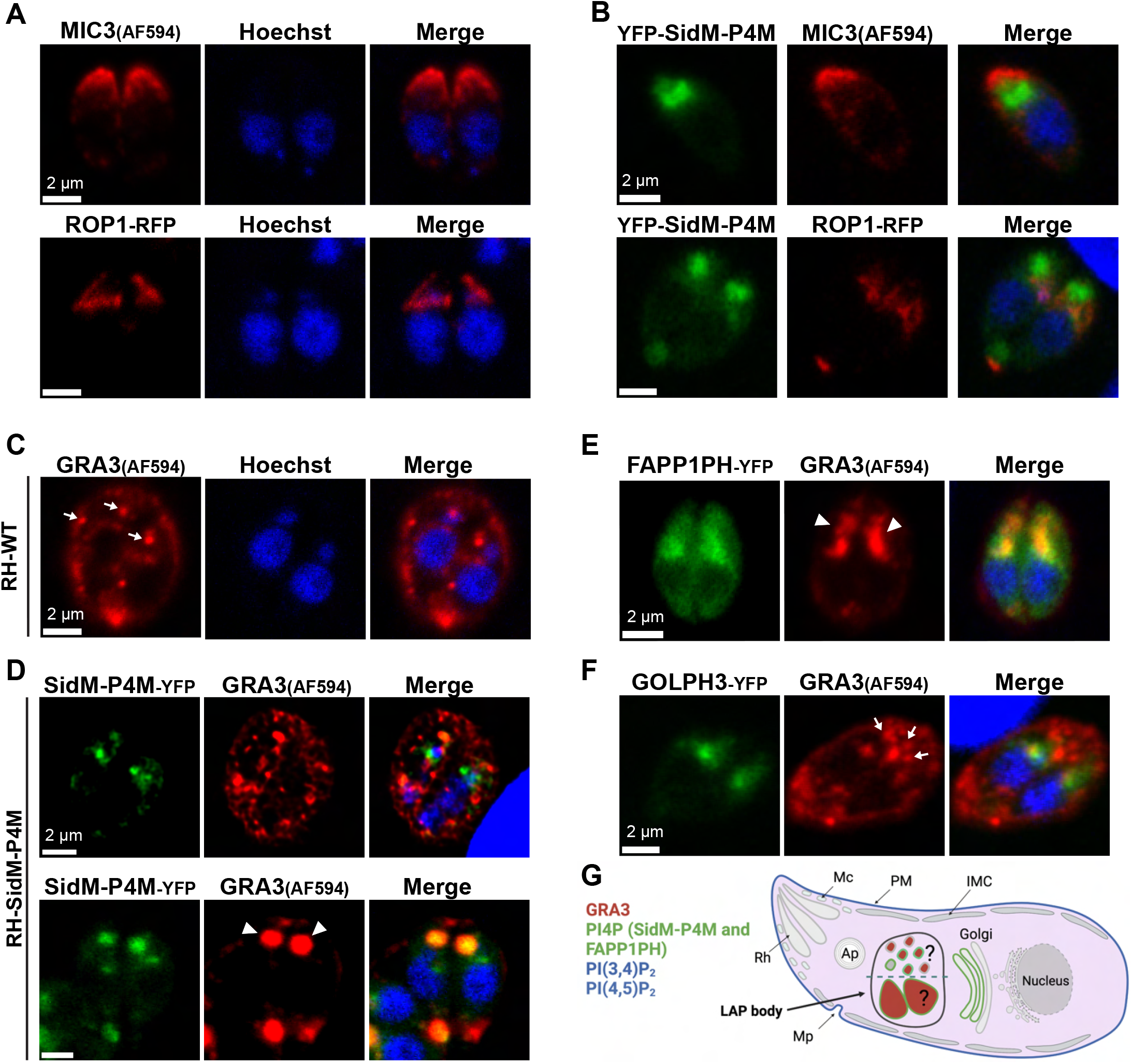
Specific derangement of DG cargo-carrying compartments in parasites expressing high affinity-PtdIns4P biosensors. Confocal microscopy images of fixed RH-WT and transgenic strains are shown. Parasite DNA is visualized using Hoechst (blue). **(A)** The microneme marker (IFA with anti-MIC3) and the rhoptry marker (ROP1-RFP) are normally distributed in RH-WT and RH-ROP1 strains, respectively. **(B)** Distribution of microneme (MIC3) and rhoptry (ROP1-RFP) markers are not affected by expression of YFP-SidM-P4M. **(C-F)** IFA of DG using GRA3 as marker (identified using anti-GRA3 imunoglobulin) during G1/S phase. **(C)** RH-WT contain scattered DGs of consistent dimensions in the cytoplasm (arrows). **(D)** Cells with a lower fluorescence signal of YFP-SidM-P4M do not form a LAP body and show WT-like GRA3 distributions (upper panel). The LAP body in RH-SidM-P4M parasites is stained with anti-GRA3 antibodies (arrowheads in lower panel). **(E)** RH-FAPP1PH parasites show a LAP body stained with anti-GRA3 antibodies (arrow heads). **(F)** DG marker distribution is typically not affected in RH-GOLPH3 strains (arrows). **(G)** Summary schematic representation of PIP pools described in this manuscript. PtdIns(4,5)P_2_ and PtdIns(3,4)P_2_ localize to the parasite plasma membrane. SidM-P4M and FAPP1PH biosensors localize PtdIns4P pools to the Golgi/TGN and the LAP body. The LAP body is only formed upon expression of these PtdIns4P biosensors and is accompanied by GRA3 localization to this structure.

By contrast, whereas immunostaining for the DG marker GRA3 in RH-WT parasites showed small DG puncta that were dispersed throughout the cytoplasm (arrows in **Figure 2C**), RH-SidM-P4M strains exhibited two distinct DG profiles that associated with the level of biosensor expression. RH-SidM-P4M parasites with low fluorescence signal, i.e. cells inferred to exhibit low PtdIns4P biosensor expression, displayed DGs with morphologies and intracellular distributions that were similar to those of WT parasites. Moreover, only an infrequent colocalization of GRA3 with the PtdIns4P biosensor was detected (Pearson’s correlation coefficient ± SEM = 0.337 ± 0.013; **Figure 2D** upper panel). The predominant population of RH-SidM-P4M parasites exhibited stronger fluorescence signals, however, and those cells were inferred to exhibit robust expression of SidM-P4M-YFP. In these parasites, strong co-localization of SidM-P4M-YFP with GRA3 was observed at the LAP body (Pearson’s correlation coefficient ± SEM = 0.893 ± 0.015; **Figure 2D** lower panel). These results were recapitulated in RH-FAPPP1H parasites (**Figure 2E**). Interestingly, the intracellular distributions and the morphologies of GRA3-positive DGs in RH-GOLPH3 parasites were similar to those of RH-WT (**Figure 2F**). However, RH-GOLPH3 parasites that exhibited GRA3 co-localization with LAP bodies with profiles similar to those of RH-SidM-P4M cells were occasionally observed. YFP-GOLPH3 expression was particularly robust in those less frequent cases. These collective results identified the LAP body as a post-Golgi compartment that houses DG cargo (**Figure 2G**), and indicated a role for PtdIns4P biosensor expression in the induction of LAP body formation.

### Expression of high-affinity PtdIns4P-binding domains induces altered intracellular distribution of multiple DG cargo

As the GOLPH3 biosensor harbors a domain with a lower PtdIns4P binding affinity than does the SidM-P4M biosensor (Wood et al., 2009; Del Campo et al., 2014), the data raised the possibility that it was sequestration of PtdIns4P from its effectors by high-affinity PtdIns4P-binding domains that induced morphological derangement of compartments involved in DG biogenesis into LAP bodies. To further evaluate and quantify LAP body formation upon sequestration of intracellular PtdIns4P pools, two genetically-encoded reporters for DG cargo were generated -- GRA3-RFP and GRA2-RFP. These constructs allowed vital monitoring of pools of DG cargo synthesized after parasite transfection and subsequent infection of host cells. Reporter constructs for each DG cargo were transiently co-transfected into RH-WT cells with an appropriate PtdIns4P biosensor construct. Live parasites were subsequently imaged. Fifty parasitophorous vacuoles (PVs) were analyzed in three independent biological replicates for each experimental condition, and DG intracellular distribution was scored as: no visible vesicles in the cytoplasm (DG-less), dispersed vesicles of typical morphology (Normal), or LAP body containing (LAP body) (**Figure 3**).

**Figure 3.**
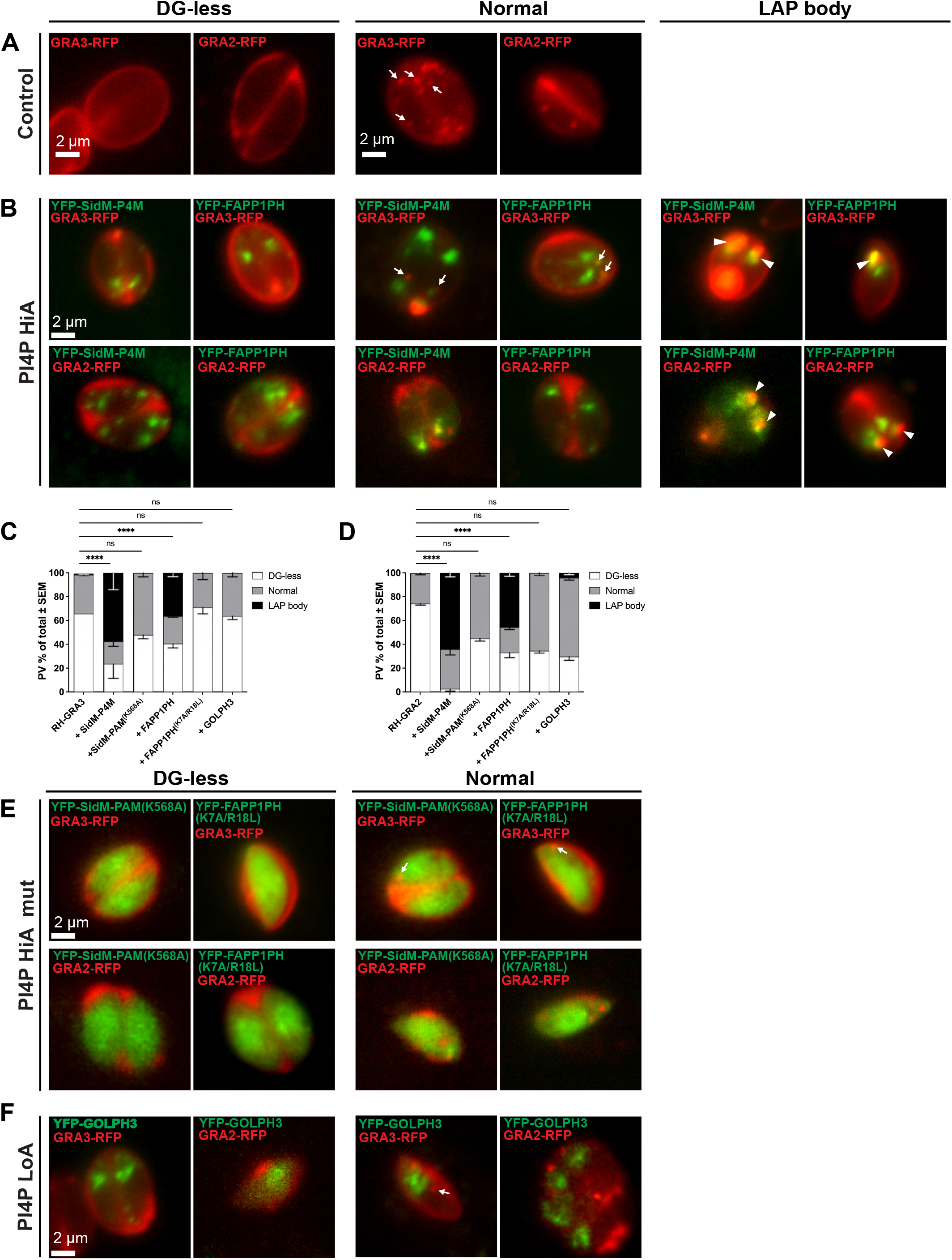
LAP body formation is induced by high affinity-PtdIns4P biosensor expression. **(A-C)** Widefield fluorescence microscopy of live intracellular parasites expressing the indicated PIP biosensors, classified by the DG reporter distribution phenotype: DG-less, normal (arrows) or LAP body (arrow heads). **(A)** Control reporter strains RH-GRA3 and RH-GRA2 exhibit either DG-less or normal cytoplasmic puncta profiles. **(B)** Parasites transiently co-expressing GRA3-RFP or GRA2-RFP in the face of high-affinity SidM-P4M- or FAPP1PH-based PtdIns4P biosensor co-expression (PI4P HiA) form a LAP body (arrow heads). **(C-D)** Quantification of PVs exhibiting the **(C)** GRA3-RFP or **(D)** GRA2-RFP phenotypes (DG-less, normal, LAP body) in the face of co-expression with the indicated biosensor (n=50 PVs). Data show mean ± SEM of three independent experiments. Statistical analyses compared the control reporter strain (RH-GRA3-RFP or RH-GRA2-RFP) and parasites co-expressing the DG reporter and the appropriate PIP biosensor with regard to LAP body phenotype. Statistical significance was calculated using two-way ANOVA followed by Dunnet’s multiple comparison test; *p* > 0.05 (ns); *p* ≤ 0.05 (*); *p* ≤ 0.01 (**); *p* ≤ 0.001 (***); *p* ≤ 0.0001 (****). **(E)** Expression of the SidM-P4M^K568A^ and FAPP1PH^K7A/R18L^ biosensor PtdIns4P-binding mutants (PI4P HiA mut) does not affect GRA3-RFP or GRA2-RFP distribution. **(F)** Parasites co-expressing the GRA3-RFP or GRA2-RFP reporter with the lower affinity (LoA) PtdIns4P YFP-GOLPH3 biosensor do not form LAP bodies.

The control condition represented by RH-GRA3 or RH-GRA2 strains presented both normal and absent DG distribution phenotypes but no LAP body formation (**Figure 3A**). In cells expressing the high affinity FAPP1PH and SidM-P4M PtdIns4P-binding domains, the DG cargo reporters redistributed into LAP bodies (**Figure 3B**, large). Quantification of those images reported a significant increase in this redistribution phenotype for both the GRA3 (mean % of PVs with LAP bodies ± SEM = 36.4% ± 3.2 for FAPP1PH and 58.5% ± 14.8 for SidM-P4M; **Figure 3C**) and the GRA2 reporters (mean % of PVs with LAP bodies ± SEM = 48% ± 5.2 for FAPP1PH and 64% ± 3 for SidM-P4M; **Figure 3D**). Thus, LAP body formation reflected altered intracellular distribution of at least two DG cargos.

That DG cargo redistribution resulted from diminished PtdIns4P signaling was demonstrated by the fact that expression of the mutant SidM-P4M^K568A^ and FAPP1PH^K7A/R18L^ biosensors induced no such effect (**Figure 3C-E**). These mutant domains are defective for PtdIns4P-binding (Del Campo et al., 2014; Jung et al., 2002) and, consistent with those defects, both mutant biosensors exhibited primarily cytoplasmic profiles (**Figure 3E**). These results demonstrate that localization of FAPP1PH- and SidM-P4M-based biosensors to the parasite Golgi/TGN system, and the morphological derangement of DG cargo-positive compartments in the form of LAP bodies, was dependent on the PtdIns4P-binding properties of these biosensors. By contrast, and consistent with DG cargo distribution representing a PtdIns4P-dependent process, expression of the lower affinity YFP-GOLPH3 biosensor failed to significantly alter the intracellular distribution of DG cargo relative to wild-type controls (**Figure 3F**).

### Altered DG morphologies report a specific PtdIns4P requirement

PtdIns4P is the metabolic precursor of the bis-phosphorylated phosphoinositides PtdIns(4,5)P_2_ and PtdIns(3,4)P_2_. To determine whether redistribution of DG cargo into LAP bodies reflected an intrinsic defect in PtdIns4P signaling, or some indirect downstream effect of PtdIns(4,5)P_2_ and/or PtdIns(3,4)P_2_ limitation under conditions of compromised PtdIns4P availability, the effects of high-affinity PtdIns4P-binding domain expression on parasite PtdIns(4,5)P_2_ and PtdIns(3,4)P_2_ pools were assessed. To that end, *T. gondii* cells transiently co-expressing FAPPPH1-RFP along with either the 2xPHPLCδ-EGFP or YFP-TAPP1PH biosensors were analyzed. Decreased production of these PIP_2_ species as a consequence of sequestration of intracellular PtdIns4P by FAPPPH1 was expected to induce release of the corresponding PIP_2_ biosensors from the plasma membrane. This prediction was not borne out by the data, however. FAPPPH1-RFP expression failed to compromise association of either PIP_2_ biosensor with the plasma membrane (**Figure 4A**).

**Figure 4.**
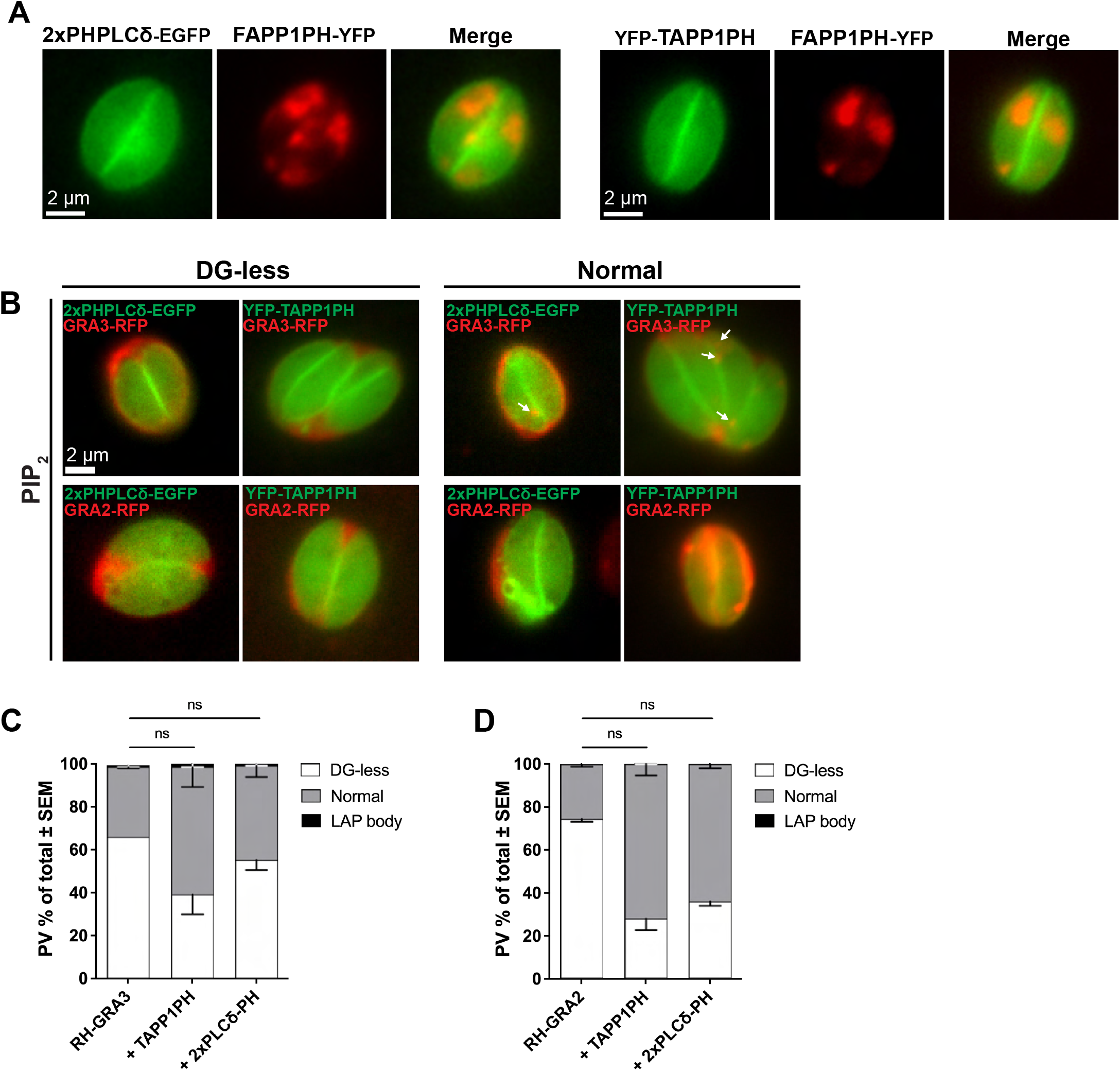
Expression of PI(4,5)P_2_ and PI(3,4)P_2_ biosensors does not induce LAP body formation. **(A)** Widefield fluorescence microscopy of live intracellular parasites co-expressing the indicated PIP biosensors: FAPP1PH-RFP and 2xPHPLCδ-EGFP or FAPP1PH-RFP and YFP-TAPP1PH. The distribution of neither PIP_2_ biosensors was affected by PtdIns4P pool sequestration at the Golgi-TGN and post-TGN compartments. **(B)** Widefield fluorescence microscopy of live intracellular parasites expressing the indicated PIP biosensors, classified by the DG reporter distribution phenotype: DG-less, normal (arrows) or LAP body (arrow heads). Expression of 2xPHPLCδ-EGFP or YFP-TAPP1PH in reporter strains RH-GRA3-RFP or RH-GRA2-RFP did not result in LAP body formation. **(C-D)** Quantification of PV phenotypes as reported by **(C)** GRA3-RFP or **(D)** GRA2-RFP cargo (DG-less, normal, LAP body) as a function of co-expression with the indicated biosensor (n=50 PVs). Data are represented as mean ± SEM of three independent biological replicates. Statistical analysis compared the control reporter strain (RH-GRA3-RFP or RH-GRA2-RFP) with parasites co-expressing the DG reporter and the appropriate PIP biosensor with regard to LAP body phenotype. Two-way ANOVA followed by Dunnet’s multiple comparison test was used to determine statistical significance; *p* > 0.05 (ns); *p* ≤ 0.05 (*); *p* ≤ 0.01 (**); *p* ≤ 0.001 (***); *p* ≤ 0.0001 (****).

As independent approach, the intracellular distributions of both the GRA2-RFP and GRA3-RFP DG cargo reporters were visualized in parasites expressing either of the two PIP_2_ biosensors. In neither case was distribution of the DG cargo reporters altered by PIP_2_ biosensor expression relative to unperturbed wild-type controls (**Figure 4B-D**). Taken together, these data report an intrinsic signaling role for PtdIns4P in the biogenesis and/or function of intracellular compartments that participate in DG cargo trafficking – an involvement distinct from the role of PtdIns4P as metabolic precursor for PtdIns(4,5)P_2_ or PtdIns(3,4)P_2_ signaling.

### Ultrastructure of DG compartments under conditions of PtdIns4P stress

To gain a more precise description of LAP body structure two independent approaches were employed. First, LAP bodies were imaged using confocal microscopy coupled with an Airyscan detector that increased spatial resolution some 1.7X and an increase in signal to noise ration of up to 8X after linear deconvolution analysis (Huff et al., 2015). This enhanced resolution is accurate down to 140 nm in the *xy*-plane – thereby allowing quantification of alterations in DG morphologies. In G1/S RH-WT parasites, GRA3-positive DGs were distributed throughout the cell (**Figure 5A**), and z-projections of confocal images indicate these DGs exhibited a mean diameter of 273 ± 7.5 nm. Interestingly, the LAP bodies of RH-YFP-SidM-P4M-expressing parasites were not simple homogeneous structures. Rather, the LAP bodies consisted of a ‘clustered’ network of smaller DGs (**Figure 5B**). These smaller DGs (sDG) exhibited a mean diameter of 140 ± 1.6 nm that was approximately half of that exhibited by the DGs of wild-type parasites (**Figure 5C**). Overlay of the GRA3 and RH-YFP-SidM-P4M profiles confirmed colocalization of these two reporters (Pearson’s correlation coefficient = 0.77 ± 0.04; black arrows in **Figure 5B**). A striking feature of the reconstructed images was that the PtdIns4P biosensor was not isotropically distributed throughout the LAP body. Rather, the biosensor, and by inference PtdIns4P, was concentrated in discrete domains throughout the sDG network (white arrows in **Figure 5B**). PtdIns4P-positive structures devoid of GRA3 were also noted -- suggesting different classes of cargo-carrying vesicles are present within the LAP body.

**Figure 5.**
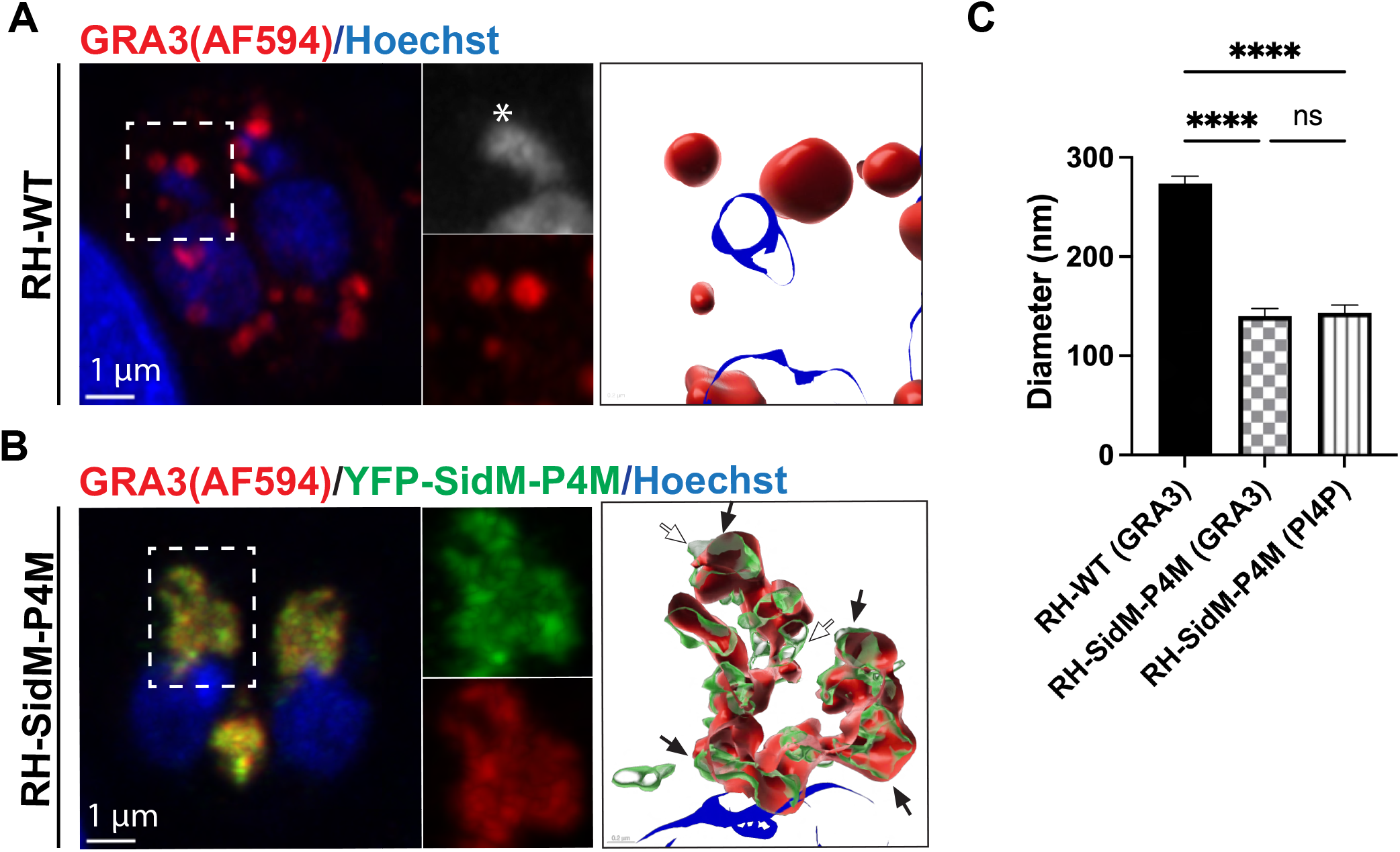
The LAP body is comprised of a network of small GRA3- and PtdIns4P-containing vesicles. **(A-B)** Z-projections of Airyscan acquisitions of RH-WT and RH-SidM-P4M parasites stained with anti-GRA3 immunoglobulin. Parasite DNA is visualized using Hoechst (blue). **(A)** Control RH-WT parasites showing DG cargo (GRA3) distributed in discrete puncta throughout the cytoplasm and PVM. G1/S phase was identified by a single undivided apicoplast in the cells of interest (asterisk). 3D-Reconstructions show the absence of a LAP body in the subapical area (dashed square). **(B)** RH-SidM-P4M parasites during G1/S phase present a LAP body that consists of numerous sDG enriched with GRA3 and PtdIns4P (dashed square). 3D-Reconstruction of the LAP body confirms spatial co-localization of GRA3 and PtdIns4P reporters (black arrows). The PtdIns4P biosensor is concentrated in distinct domains (white arrows). **(C)** Quantification of the diameter (nm) of sDGs in RH-WT parasites (control) and sDGs of RH-SidM-P4M parasites (n = 11 for both cases). Mean values ± SEM are shown. Statistical analyses used one-way ANOVA followed by Tukey’s multiple comparison test to assess significance; *p* > 0.05 (ns); *p* ≤ 0.05 (*); *p* ≤ 0.01 (**); *p* ≤ 0.001 (***); *p* ≤ 0.0001 (****).

As an independent means of probing LAP body structure with higher precision, we employed correlative light electron microscopy (CLEM). Electron microscopy analysis required confident and unbiased identification of YFP-SidM-P4M-expressing parasites in the electron microscopy grid -- a requirement made difficult by: (i) the low efficiency with which the biosensor construct is transfected into parasites, and (ii) our inability to create stable expression cell lines for the high affinity PtdIns4P biosensors. As such, CLEM was employed to specifically target YFP-SidM-P4M-expressing parasites for further EM imaging. To that end, human foreskin fibroblasts were seeded onto gridded glass-bottom coverslip dishes, infected with RH-SidM-P4M parasites, and fixed parasites were stained with antibodies directed against GRA3. The positions of either RH-WT or RH-SidM-P4M parasites were registered in the coordinate system of the gridded coverslip by confocal microscopy. RH-SidM-P4M parasites in which LAP bodies were evident by confocal microscopy and identified as regions of interest (ROI) and specifically selected for CLEM imaging. The grids were then processed for transmission EM (TEM) and the ROIs visualized by transmission electron microscopy.

In excellent agreement with the results of the Airyscan imaging experiments, the TEM acquisitions reported well dispersed mature DGs (**Figure 6A**). Again, consistent with the Airyscan imaging data, RH-SidM-P4M-expressing parasites presented the LAP body as a single large cluster of electron dense vesicles in the apical region of the cell (green arrows in **Figure 6B**). The electron dense vesicles in RH-SidM-P4M were approximately 5X more numerous than the DGs of wild-type parasites (mean sDG count per EM section ± SEM = 3 ± 0.5 for RH-WT and 14 ± 3.7 for SidM-P4M; **Figure 6C**), and were also about half the dimension of those observed in RH-WT parasites (mean DG diameter ± SEM = 242 ± 8.7 nm for RH-WT and 145 ± 4.7 nm for SidM-P4M; **Figure 6D**). Morphometric values for DGs obtained in the wild-type strain are in excellent agreement with previous measurements (Dubey et al., 1998). CLEM analyses of RH-FAPP1PH parasites yielded comparable results and, in this case, the reduced electron densities of the ‘clustered’ sDGs that comprise the LAP body were particularly noteworthy (**Figure 6E**). Thus, data from two independent high resolution imaging approaches converge on the conclusion that the LAP body formed upon parasite expression of high affinity PtdIns4P binding domains represented a ‘clustered’ network of small vesicles that carry DG cargo.

**Figure 6.**
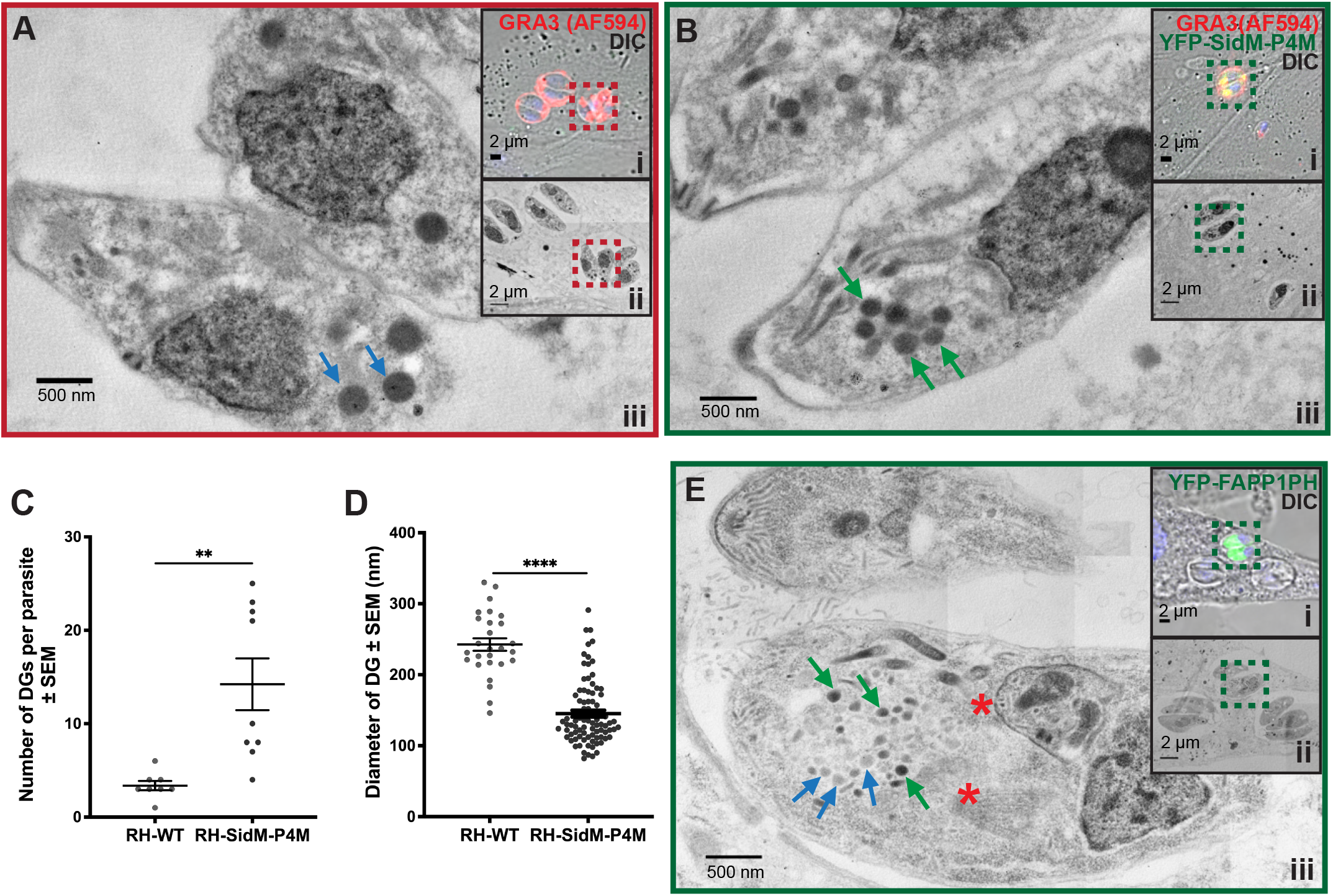
High resolution imaging of LAP body structure by correlative light electron microscopy. **(A, B and E)**. CLEM acquisitions of stained RH-WT or transgenic parasites involved the following steps: (i) targeting and recognition of ROIs (dashed square) was performed in a confocal microscope and selected for imaging by TEM at (ii) low and (iii) high magnification. **(A)** RH-WT show scattered distribution of mature highly electro-dense DGs distributed throughout the cytoplasm (red arrows). **(B)** RH-SidM-P4M show a pool of DG-cargo carrying vesicles in the subapical area that are of smaller size (sDGs) relative to WT DGs (green arrows). **(C)** Quantification of DG number per tachyzoite and per EM section (n = 8 for WT and SidM-P4M). Data represent the mean ± SEM. **(D)** Quantification of the average DG diameter (nm) per tachyzoite and per EM section (n =8 for WT and SidM-P4M and n=3 for FAPP1PH). Data are represented as mean ± SEM, and statistical significance was determined using the Mann-Whitney t-test; *p* > 0.05 (ns); *p* ≤ 0.05 (*); *p* ≤ 0.01 (**); *p* ≤ 0.001 (***); *p* ≤ 0.0001 (****). **(E)** An RH-FAPP1PH parasite undergoing endodyogeny (red asterisks indicate position of daughter cells) presents a LAP body consisting of an ‘clustered’ sDG network (green arrows) that resides in the subapical region of the mother cell. sDGs with reduced electron density are indicated by blue arrows.

### PtdIns4P stress compromises DG cargo exocytosis

Deranged organelle morphology is often associated with compromised function. Thus, we tested whether expression of high-affinity PtdIns4P-binding domains compromised exocytosis of DG cargo. To quantify exocytosis, RH-WT and RH-SidM-P4M parasites were immunostained for GRA3 and the intensities of exocytosed GRA3 signal in the PV membrane (PVM) were ratioed to total GRA3 intensity in the PV as a measure of DG exocytosis efficiency. For RH-SidM-P4M cells, only parasites with GRA3-containing LAP bodies were analyzed. Acquisition parameters were normalized across all samples to minimize technical bias.

The data show a ca. 58% reduction in normalized GRA3 signal at the PVM of SidM-P4M-dependent LAP body-containing parasites in comparison with RH-WT (**Figure 7A and B**). That the observed DG exocytic defects were the consequences of reduced PtdIns4P signaling was indicated by the fact that the PVM GRA3/total GRA3 ratios measured for parasites expressing the PtdIns4P-binding mutant of SidM-P4M (RH-SidM-P4M^K568A^ parasites) were not significantly diminished compared to those of RH-WT (**Figure 7B**). Taken together, these collective data demonstrate that PtdIns4P stress in *T. gondii* results in: (i) altered DG biogenesis, and (ii) compromise of the normally efficient exocytosis of DG contents.

**Figure 7.**
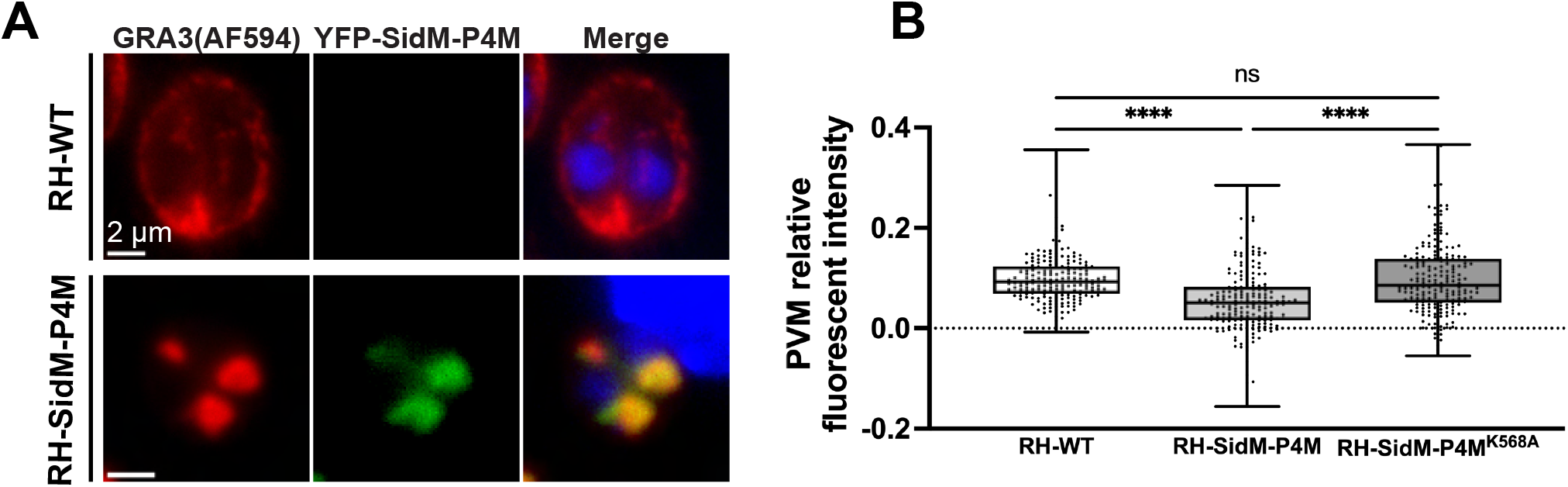
Sequestration of PtdIns4P inhibits secretion of GRA3 to the PVM. **(A)** Widefield fluorescence microscopy of fixed intracellular parasites immunostained with anti-GRA3 immunoglobulin in the absence (upper panel) or presence (lower panel) of SidM-P4M-YFP expression. In RH-WT cells, GRA3 localizes to the PV and PVM. In RH-SidM-P4M cells, GRA3 is mainly retained in the LAP body in the intracellular apical region and is absent from the PVM. Parasite DNA is visualized using Hoechst (blue). **(B)** Quantification of fluorescence intensity of GRA3 in the PVMs of parasites exhibiting SidM-P4M-dependent LAP body formation is significantly reduced compared to that of PVMs of RH-WT and SidM-P4M^K568A^– expressing strains. The box and whisker plot shows the average of indicated individual measurements ± SEM per experimental condition. Individual measurements were collected from two independent biological replicates (n=196). Statistical significance was calculated using one-way ANOVA followed by Tukey’s multiple comparison test; *p* > 0.05 (ns); *p* ≤ 0.05 (*); *p* ≤ 0.01 (**); *p* ≤ 0.001 (***); *p* ≤ 0.0001 (****).

### PtdIns4P biosensor distribution through the parasite cell cycle

PtdIns4P distribution profiles were also followed throughout the parasite cell division cycle. For those experiments, an inner membrane complex (IMC) reporter was constructed where the IMC marker IMC1 was fused to RFP (IMC1-RFP) to landmark cell cycle stage in proliferating intracellular parasites – as has been done previously (Ouologuem and Ross, 2014). Live parasites expressing IMC1-RFP and YFP-SidM-P4M were imaged by widefield fluorescence microscopy during various stages of the cell cycle. Representative images collected for each cell cycle stage are shown in **Figure 8A**. In G1/S parasites, YFP-SidM-P4M localized primarily to the LAP body in agreement with the various data documented above. At initiation of cell division, a prominent recruitment of YFP-SidM-P4M to a split juxta-nuclear stacked compartment was observed (white arrows in **Figure 8Ai**) and, to a lesser extent, to apical vesicles (yellow arrows in **Figure 8Ai**). Quantification of parasites with YFP-SidM-P4M distribution to the LAP body demonstrated that, contrary to what was observed in G1/S parasites or for cells in the next division stage (elongation 1), only ∼8.5% of cells at the initiation phase exhibited positive signal at the LAP body. The remaining cells exhibited profiles that reported redistribution of YFP-SidM-P4M to the split juxta-nuclear stacked compartment (**Figure 8Aii and iii**). During cell elongation stages, that stacked compartment completed fission and each daughter cell inherited a copy. Thus, the split juxta-nuclear compartment exhibited the features expected of a dividing Golgi system (Pelletier et al., 2002). That this stacked compartment indeed represented the Golgi/TGN was confirmed by the fact that it was marked with TgSORTLR (**Figure 8B**).

**Figure 8.**
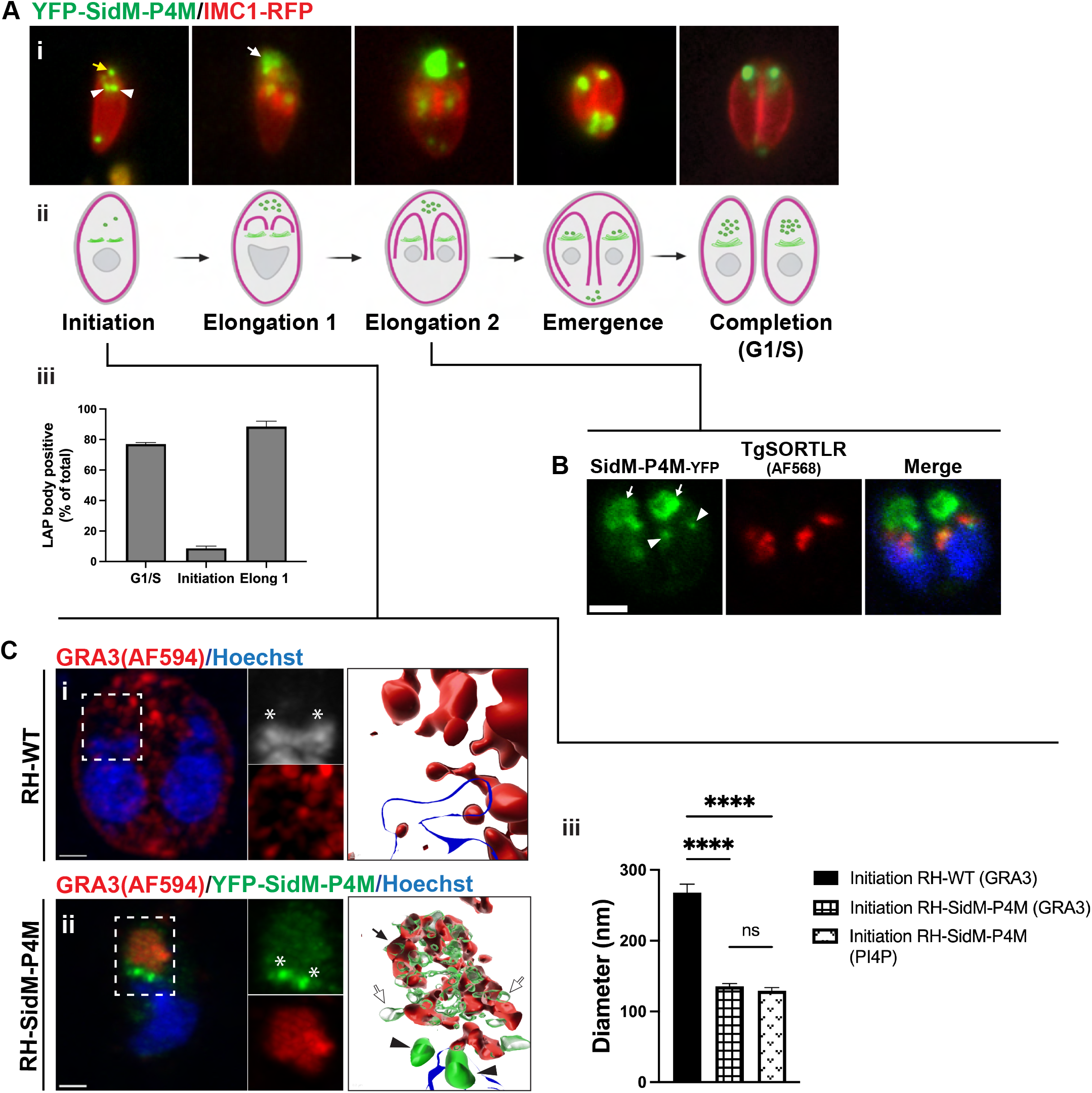
PtdIns4P biosensor relocates away from the LAP body during the initial stages of parasite cell division. **(Ai)** Widefield fluorescence microscopy images of live cells co-expressing YFP-SidM-P4M and cell cycle marker IMC1-RFP at each step of the cell cycle are shown. A dividing Golgi system marks initiation of endodyogeny (arrowheads). YFP-SidM-P4M targets to the LAP body during most stages of the cell cycle (white arrows). During the initiation stage, however, YFP-SidM-P4M redistributes to the dividing Golgi system (arrowheads) and to apical vesicles (yellow arrow). **(Aii)** Schematic representation of the relative positions of the IMC (magenta) and of PtdIns4P pools (green) at the Golgi/TGN and post-TGN compartment at each cell cycle stage (initiation, elongation (1 and 2), emergence and completion/G1). **(Aiii)** Quantification of parasites showing recruitment of YFP-SidM-P4M to the LAP body at the indicated stages of endodyogeny. Quantification data were obtained from two independent experiments (n ≤ 15 parasites per stage). Data show mean ± SEM. **(B)** Confocal IFA images identifying Golgi/TGN (TgSORTLR) in a dividing RH-SidM-P4M parasite during the elongation stage of cell division. YFP-SidM-P4M targets to the dividing Golgi/TGN compartment (arrowheads) and the LAP body (arrows). DNA is visualized using DAPI (blue). **(C)** Z-projection of Airyscan acquisitions of parasites immunostained with anti-GRA3 immunoglobulin during initiation of endodyogeny. This stage is identified by the division of apicoplast in RH-WT (DNA visualized with Hoechst; asterisks) and fission of the Golgi/TGN system in RH-YFP-SidM-P4M cells (asterisks). **(Ci)** Control RH-WT parasites show GRA3 distributed in discrete puncta throughout the cytoplasm and PVM, while the RH-YFP-SidM-P4M strain displays a GRA3 positive-LAP body with low YFP-SidM-P4M signal **(Cii)**. Rather, YFP-SidM-P4M targets to the recently divided Golgi/TGN. 3D reconstruction demonstrates two regions within the LAP body: GRA3+/PtdIns4P+ (black arrows), and GRA3-/PtdIns4P+ (white arrows). Parasite DNA is visualized using Hoechst (blue). **(Ciii)** Quantification of the sDG diameter (nm) in dividing RH-WT (control) vs RH-YFP-SidM-P4M parasites (n = 11 for both cases). Data are presented as mean ± SEM, and statistical significance was determined using one-way ANOVA followed by Tukey’s multiple comparison test; *p* > 0.05 (ns); *p* ≤ 0.05 (*); *p* ≤ 0.01 (**); *p* ≤ 0.001 (***); *p* ≤ 0.0001 (****).

By contrast to the Golgi system, the LAP body was neither divided nor inherited. At elongation stages of the cell cycle, YFP-SidM-P4M partially re-localized back to the LAP body of the mother cell and, upon daughter cell emergence, the LAP body migrated towards the basal pole of mother cell body remnants --potentially to the residual body. At completion of the cell cycle (marked by entrance of daughter cells into G1), YFP-SidM-P4M redistributed primarily to a LAP body profile. Thus, the biosensor data report a mobilization of the PtdIns4P biosensor from the LAP body to a newly dividing Golgi system upon initiation of endodyogeny, and relocalization of the biosensor back to the LAP body in subsequent stages of the cell cycle.

### PtdIns4P biosensor redistributes away from the LAP body in dividing parasites

The apparent ‘disappearance’ of the PtdIns4P-positive LAP body in parasites initiating cell division, suggested two possibilities: (i) the LAP body is disassembled during initiation of endodyogeny and subsequently reforms in later stages of the cell cycle, or (ii) the biosensor relocates away from an otherwise intact LAP body. To distinguish between these possibilities, LAP body structure was examined in dividing parasites by high resolution Airyscan confocal imaging with the modification that GRA3 replaced YFP-SidM-P4M as LAP body marker. In RH-WT parasites, GRA3-positive DGs were distributed throughout the cell during the initiation of mitosis as landmarked by the elongation of the apicoplast (asterisk in **Figure 8Ci**; Martins-Duarte et al., 2021). By contrast, parasites expressing YFP-SidM-P4M during the initiation of cell division exhibited a GRA3-positive LAP body (**Figure 8Cii**). Whereas the RH-WT DGs exhibited a mean diameter of 268 ± 11.5 nm – a value consistent with that measured for DGs of RH-WT cells in G1/S phase (273 ± 7.5 nm; see above) -- the sDGs that comprised the LAP body in those dividing parasites presented dimensions similar to those of YFP-SidM-P4M-expressing parasites in G1/S phase (mean ± SEM in G1/S phase: 140 ± 7.6 nm vs in mitosis: 135 ± 3.8 nm) (**Figure 8Ciii**). This GRA3-positive structure remained prominent even though SidM-P4M PtdIns4P biosensor localization to the LAP body was strongly diminished during the initiation of cell division (Pearson’s correlation coefficient 0.39 ± 0.05; white arrows in **Figure 8Cii**, compared to 0.77 ± 0.04 for G1/S phase cells). The redistribution of YFP-SidM-P4M from the LAP body to the Golgi/TGN system at the initiation of cell division, and its restored targeting to the LAP body in subsequent stages of the cell cycle, suggests a complex and previously unappreciated regulation of PtdIns4P signaling in DG cargo-containing compartments upon initiation of parasite cell division.

## DISCUSSION

Current models envision formation of DGs to result via direct budding from the *T. gondii* TGN as mature structures with appropriately sorted and condensed cargo (Griffith et al., 2022). Thus, DGs are not considered to undergo the types of more complex maturation processes that are hallmarks of dense core vesicle compartments that form the basis of regulated exocytosis systems of other eukaryotes – e.g. dense core granules of neuroendocrine cells or insulin secretory granules (ISGs) of pancreatic β-cells. Direct budding models are based on imaging studies that document the presence of morphologically uniform mature DGs with no recognizable maturation intermediates (Karsten et al., 1998; Coppens et al., 1999). Direct budding mechanisms leave unanswered questions, however. Although there is considerable evidence to suggest that the DG exocytic pathway is a default pathway in *T. gondii* (Heaslip et al., 2016; Striepen et al., 1998), these models nevertheless require that DG cargo be segregated from other secretory cargo and condensed – presumably at the site of DG budding from the TGN surface. Moreover, proteins marked for retention in the TGN must either be excluded from loading into the nascent DG vesicle or otherwise retrieved from a DG vesicle after scission from TGN membranes. How appropriate cargo is condensed into these electron dense granules remains an unresolved question in the field of apicomplexan secretory trafficking. Mechanisms of cargo quality control are also not understood in detail. These questions are of particular interest as efficient trafficking and exocytosis of virulence factors is fundamental to host cell invasion by the parasite, its intracellular proliferation, and ultimate egress of the parasite from the exhausted host cell to initiate successive rounds of infection.

### Reconsidering ideas for DG biogenesis

While existing imaging data argue for direct DG budding models based on the lack of morphological evidence for discrete steps in the DG biogenic process, our findings indicate that DG formation in *T. gondii* is a more complex process than that envisioned by simple direct budding models – one perhaps more in line with the pathways of dense core vesicle and secretory granule biogenesis in other systems. Using neuroendocrine cells as example, dense core granules mature in a stage-specific progression of homotypic fusion and retrograde trafficking of missorted cargo back to the TGN (Merighi, 2018). Moreover, the maturation process is typically slow in systems where immature DG-like structures are visible under the electron microscope. For example, ISG biogenesis in mammalian pancreatic β-cells is not only a high flux pathway, but ISG maturation requires three hours to complete (Davidson et al., 1988; Omar-Hmeadi et al., 2021). The lack of information regarding discrete stages of DG biogenesis in *T. gondii* is potentially a reflection of the process being too rapid and/or of insufficient flux to permit confident visualization of transient biogenic intermediates by electron microscopy – i.e. the only method with the resolution necessary to recognize such intermediates.

As demonstrated in other systems (Zhang et al., 2017; Tandon et al., 1998), either in vitro reconstitution or in vivo stage-specific inhibition approaches are required to systematically dissect DG biogenesis to arrive at a detailed description of the process in *T. gondii*. We argue that it is from the latter perspective that the expression of high affinity PtdIns4P protein domains in *T. gondii* is particularly informative. Our description of the LAP body as a ‘clustered’ network of small PtdIns4P-decorated vesicles loaded with DG cargo argues for reconsideration of direct budding models for DG biogenesis in *T. gondii*. Furthermore, the fact that LAP body formation is induced by high affinity PtdIns4P biosensor expression demonstrates an important and previously unappreciated role for PtdIns4P signaling in DG biogenesis. Finally, the morphological properties of the LAP body itself offer other insights into this process. In **Figure 9**, we outline a hypothetical model for DG biogenesis that takes these insights into account.

**Figure 9.**
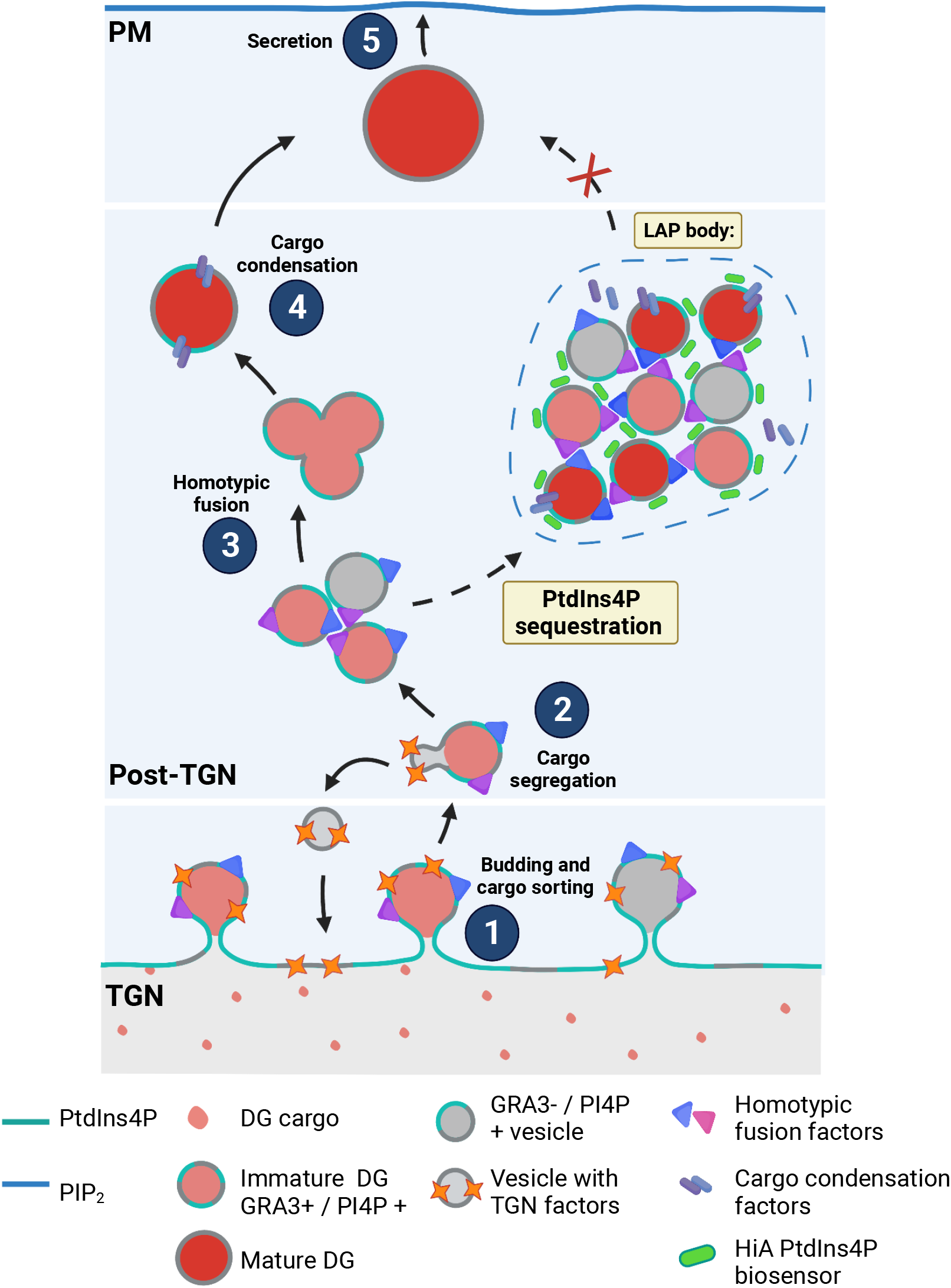
Model for DG biogenesis, maturation and exocytosis. Solid arrows mark proposed model for DG biogenesis and maturation in *T. gondii*. Budding and cargo sorting from PtdIns4P-enriched domains of the TGN (**step 1**) are followed by a further round(s) of cargo segregation (**step 2**) to generate immature DGs. Maturation of these structures at a post-TGN compartment is characterized by the PtdIns4P-dependent homotypic fusion (**step 3**) – interference of which results in LAP body formation. Subsequent steps involve cargo condensation in the maturing DGs (**step 4**). Removal of PtdIns4P from mature DGs membrane marks completion of the maturation process and acquisition of secretion competence (**step 5**).

### Functional interpretation of the LAP body

How do we interpret the LAP body and how does this structure relate to DG biogenesis? We favor the idea that the LAP body represents an exaggerated compartment of nascent DGs arrested at a late stage(s) of biogenesis that we loosely refer to as maturation. The morphological data indicate that the small DG cargo-containing vesicles that comprise the LAP body network have completed budding from the TGN (**Figure 9**, step 1). Those data further suggest these have largely completed the process of cargo segregation (**Figure 9**, step 2).

These conclusions are supported by demonstrations that the small LAP body vesicles are loaded with multiple DG cargo, are of homogeneous dimensions and are devoid of detectable quantities of TGN resident proteins (e.g. TgSORTLR). These data demonstrate the small vesicles are not a result of TGN fragmentation induced by expression of high affinity PtdIns4P binding modules. The fact that these vesicles are smaller and (in some circumstances) less electron dense than the mature DGs of WT parasites suggests a late maturation step(s) is perturbed. In turn, these perturbations compromise exocytosis of DG cargo.

### A role for PtdIns4P in DG biogenesis/maturation

As LAP body formation requires expression of high affinity PtdIns4P-binding modules, we interpret LAP body formation as a consequence of biologically insufficient PtdIns4P signaling in the process of DG biogenesis/maturation. The small dimensions of the clustered LAP body vesicles relative to mature DGs of WT cells suggests that homotypic fusion of the small clustered DGs is a discrete PtdIns4P-requiring step in the formation of the larger mature DGs (**Figure 9**, step 3). That PtdIns4P distribution within the LAP body and on the small vesicle surfaces is not isotropic suggests the existence of discrete PtdIns4P domains on clustered DGs. It is tempting to speculate that these domains might be concentrated in vesicle factors required for homotypic tethering and/or fusion. In that regard, mammalian COPII coated vesicles derived from the endoplasmic reticulum engage in homotypic tethering to form vesicular/tubular clusters (VTCs; Yu et al., 2006). Although not known to be a PtdIns4P-dependent process in the case of COPII vesicles, the general morphological similarities in the VTCs and LAP bodies are notable. Alternatively, given that the clustered LAP body DGs are in some circumstances less electron dense than the mature DGs of WT parasites, PtdIns4P might also be required for the optimal activity of factors required for cargo condensation (**Figure 9**, step 4). For example, cargo condensation in dense core granules of other systems require recruitment of proton pumps into the maturing DG to acidify the compartment and potentiate cargo packing into crystalline forms (Rhodes et al., 1987; Schoonderwoert et al., 2000). The existence of PtdIns4P-positive structures in the LAP body devoid of GRA3 cargo suggest different classes of vesicles (as defined by cargo) are present in the network. Fusion of immature DGs with these vesicles might contribute to delivery of factors that regulate DG cargo condensation.

### Signals for completion of DG maturation and competence for exocytosis

Models positing a Ptdins4P signaling involvement in DG maturation must account for how completion of maturation is recognized. In that regard, LAP-body DGs are marked with both PtdIns4P and DG cargo, whereas mature DGs do not recruit high affinity PtdIns4P biosensors. We infer from those data that mature DGs are either devoid of, or present very low amounts of, PtdIns4P on their surfaces. These results raise the intriguing possibility that loss of PtdIns4P from the cytosolic face of the DG membrane marks completion of the maturation process and acquisition of competence for exocytosis (**Figure 9**, step 5). Such a strategy applies to the maturation pathway of constitutive post-Golgi secretory vesicles in yeast where the PtdIns4P loaded onto the nascent vesicle during the budding process is degraded as secretory vesicles approach sites of exocytosis – a lipid remodeling event that reprograms the small GTPase specificity of vesicle-associated nucleotide exchange factors (Ling et al., 2014). PtdIns4P pools on ISG surfaces must also be degraded by the PtdIns4P-phosphatase Sac2 prior to mature granule association with the plasma membrane (Olsen et al., 2003, Nguyen et al., 2019).

With regard to regulation of PtdIns4P levels, we observed that high affinity PtdIns4P biosensors significantly redistributed away from the LAP body to the dividing Golgi system in parasites initiating endodyogeny. The simplest interpretation of those results is that LAP body PtdIns4P levels are reduced during initiation of endodyogeny and that the biosensors return to what is typically a PtdIns4P-enriched compartment. Such remodeling might reflect a shift in emphasis in specificity of cargo exocytosis at this cell cycle stage, or perhaps PtdIns4P levels increase in the Golgi system at this stage to promote division of this compartment. In any event, as the reduction of PtdIns4P on the surfaces of LAP body vesicles was not sufficient to induce their exocytosis, we interpret the LAP body as an exocytic ‘backwater’ that is not subject to efficient chase to an exocytosis-competent intermediate. Perhaps this is why the structure migrates towards the residual body of the mother cell during endodyogeny.

### Primordial functions for PtdIns4P signaling

Apicomplexa are ancient eukaryotes that serve as excellent model organisms for the study of root mechanisms of eukaryotic cell function and evolution. In that regard, the collective data identify a role for Ptdins4P in the maturation of DGs and ultimate exocytosis of DG cargo in *T. gondii*. The similarities to aspects of both constitutive and regulatory secretory pathways in other eukaryotes are noteworthy as described above. This is perhaps not surprising as *T. gondii*, given its parasitic lifestyle, can be considered a professional secretory cell – one for which there is evidence for both constitutive and regulated DG secretory pathways (Chaturvedi et al, 1999; Coppens et al., 1999; Griffith et al., 2022).

The collective data now raise key questions regarding: (i) which PtdIns 4-OH kinase(s) produces the PtdIns4P pool(s) whose sequestration induces LAP body formation and defects in DG exocytosis, and (ii) how is synthesis of this PtdIns4P pool(s) regulated? With regard to the latter point, the LAP body phenotypes largely recapitulate those recently described in mammalian pancreatic β-cells where various stages of ISG formation, maturation and regulated exocytosis are PtdIns4P-dependent processes whose execution is perturbed upon functional ablation of PITPα – a PtdIns/phosphatidylcholine transfer protein (PITP) that stimulates PtdIns4P production in pancreatic β-cell TGN membranes (Yeh et al., 2022). Similarly, the major yeast PITP Sec14 plays an essential role in potentiating PtdIns4P-dependent membrane trafficking through the yeast TGN/endosomal system (Bankaitis et al., 1990; Hama et al., 1999; Rivas et al., 1999; Schaaf et al., 2008; Bankaitis et al., 2010; Grabon et al., 2019).

Does *T. gondii* utilize a PITP in a similar manner for PtdIns4P-dependent DG activities? This seems probable as *T. gondii* potentially encodes ten Sec14-like PITPs and one mammalian PITPα-like ortholog (ToxoDB (v56); http://ToxoDB.org). Moreover, time-resolved phosphoproteome analyses identified a Sec14-like protein (TGME49_254390) and a putative PtdIns 4-OH kinase (TGME49_276170) in a cohort of lipid signaling proteins phosphorylated upon induction of egress -- a process dependent on phosphatidic acid and phospholipase C signaling (Herneisen et al., 2022, Nofal et al. 2022). Yet another of the *T. gondii* Sec14-like PITPs (TGME49_213790) is appended to a PH domain -- as is the single PITPα-like ortholog (TGME49_289570). Such domain architectures recommend these putative PITPs as attractive candidates for engaging the PtdIns4P pool(s) identified herein. Functional dissection of these activities promises to add additional layers of complexity to the mechanism of how PtdIns4P signaling regulates DG biogenesis, maturation and exocytosis in apicomplexan parasites.

## MATERIALS AND METHODS

### Cell culture and *T. gondii* strains

The human foreskin fibroblast-1 (HFF-1) cell line was obtained from the American Type Culture Collection and grown in Dulbecco’s modified Eagle’s medium with 10% fetal bovine serum (FBS) and incubated at 37°C with 5% CO_2_ in a humidified atmosphere. The *T. gondii* RH strain was maintained by serial passage using HFF-1 monolayers as host cells. Parasites were cultured to egress and purified from host cells by disrupting the host cells via serial passage through 18- and 25-guage needles. Parasites were subsequently collected from the host cell lysate in the pellet after centrifugation of the crude lysate at 900 x g for 5 minutes.

### Generation and selection of stable transgenic parasites

RH strain was transfected by electroporation as described previously (Soldati and Boothroyd, 1993). Using an electroporation cuvette as vessel, 10^7^ tachyzoites were mixed with 25 μg of plasmid in transfection buffer (120mM KCl, 150μM CaCl_2_, 5mM MgCl_2_, 2mM EDTA, 25mM HEPES KOH, 10mM potassium phosphate, pH 7.6). In co-transfection experiments, 12.5 μg of each plasmid was used. A single electrical pulse (1.3kV, 25uF) was applied using the Bio-Rad Gene Pulse II electroporation apparatus. Parasites were allowed to recover after plating on an HFF-1 monolayer in the absence of drug for 24 hours. When appropriate, selection of stable transgenic parasites was carried out in the presence of a chloramphenicol (20 μM) selection. Stable clones were isolated by limiting dilution under conditions of continuous drug selection.

### Plasmid constructs

All *T. gondii* expression vectors expressing fluorescently-labeled organelle markers were based on the pTUB-CAT (Chloramphenicol acetyltransferase) system (Kim et al., 1993). Gene expression is driven by the *tubulin* promoter. In all cases, the DNA restriction fragments carrying the gene of interest were subcloned into the BglII/AvrII sites of pTUB. These constructs were engineered such that these carried a Myc-tag and a fluorescent protein gene between flanking the cloning site. Additional details regarding constructs of interest are described in the Figure legends. The primers used for cloning all *T. gondii* biomarkers are listed in **Supplementary Table 1**.

### Immunofluorescence analysis (IFAs)

Confluent HFF-1 host cells grown in 35mm glass-bottom coverslip dishes were infected with tachyzoites for 24 hrs, washed 1X with phosphate buffered saline (PBS) 1X and then fixed with 4% PFA (v/v) for 15 min at room temperature and washed 3X (all washing incubations were performed for 10 min with PBS 1X solution). Cells were permeabilized with 0.2% Triton-X (v/v) for 4 min at RT, blocked with 2% BSA in PBS 1X overnight at 4°C, incubated with organelle-specific primary antibodies diluted in blocking buffer for 1 hour. Details regarding the primary and secondary antibodies used in this study are listed in **Supplementary Table 2**. Coverslip were washed with PBS three times, incubated with conjugated secondary antibodies listed in **Supplementary Table 2** in blocking buffer for 1 hour, and washed again with PBS three times. Cell DNA was stained with Hoechst solution (1:5000) for 5 min. After three washes with PBS, coverslips were flooded with PBS and stored at 4&C for no longer than seven days.

### Image acquisition

Confocal images were collected in a NikonA1R microscope (CFI Plan Apo lambda 60x/1.4 oil objective) and Zeiss LSM 780 NLO multiphoton microscope (Plan-Apochromat 63x/1.4 oil DIC M27 objective). Imaging processing was performed using ImageJ (FIJI) (NIH). Pearson’s correlation coefficients were calculated using the colocalization analysis plugin JaCoP in FIJI (ImageJ) and with the colocalization analysis tool in ZEN Blue 2.3. Super-resolution fluorescence imaging was performed with a Zeiss LSM 780 NLO multiphoton microscope equipped with an Airyscan detector (Plan-Apochromat 63x/1.4 oil DIC M27 objective). Images obtained with Airyscan detector were deconvoluted using ZEN 2.3 (blue edition) software, and gamma values were increased or decreased in the order of 0.1-0.8 units). Pearson’s correlation and colocalization coefficients of regions of interest (subapical areas of individual tachyzoites) were calculated using ZEN blue software. 3D reconstructions were generated using the 3D view module in Imaris 9.8v.

### Definition and quantification of GRA3 profiles

Transgenic parasites were generated by electroporation as described above. Two plasmid pTUB constructs were used for co-transfection: 12.5 μg of pTUB-Myc-GRA3-RFP and 12.5 μg of pTUB constructs expressing YFP-FAPP1PH, YFP-SidM-P4M, YFP-GOLPH, YFP-SidM- P4M^(K568A)^, FAPP1PH^(K7A/R18L)^, 2xPHPLCδ-EGFP, or TAPP1PH-YFP as appropriate. Transfected tachyzoites were plated onto confluent HFF-1 cells in 35mm glass-bottom coverslip dishes with 2 ml of supplemented DMEM. Plates were incubated for 24 hours. Only parasites expressing the constructs of interest were counted using an epifluorescence microscope (Nikon Eclipse T*i*).

The criteria used for binning parasite populations based on GRA3 and GRA2 phenotype was as follows: 1) Each unit was considered as a PV that contained one or more tachyzoites; 2) up to 50 PVs were counted per experiment; 3) PVs were binned as displaying large or small puncta phenotype when at least one of the tachyzoites in the vacuole presented one of these phenotypes. In cases where both small and large puncta pattern were observed in the same PV, the PV was classified as displaying large puncta. All experimental results represent the data from three independent biological replicates.

### Ratiometric measurement of DG exocytosis efficiency

Wild-type and transiently transfected parasites were grown for 24 h, fixed with 4% PFA and stained with anti-GRA3 as mentioned previously. Imaging and counting of 100 PVs per experimental and control conditions was performed in epifluorescence mode. Camera settings such as format (no binning), exposure time (10 msec) and dynamic range (11-bit and Gain 4) were fixed for the TRITC filter channel to collect unbiased imaging data for all experimental conditions. To calculate GRA3 secretion efficiency as a function of transient YFP-SidM-P4M expression, we adopted the ratiometric approach of Li et al (2022). Regions of interests (ROI) were drawn around the PVM region (PVM), the intracellular region occupied by the nucleus (nuclear region), the area surrounding the entire PV and parasites (Total Toxoplasma), and a background region outside the PV. The mean intensities for each region were measured in ImageJ (FIJI) and the PVM relative intensity was obtained as:

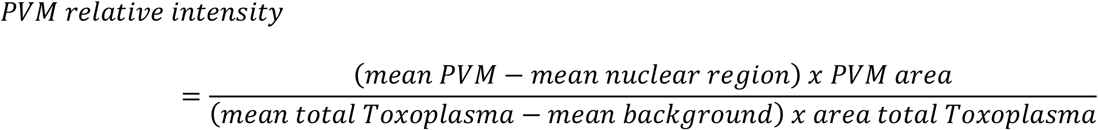

### Correlative light electron microscopy

Mattek glass-bottom coverslip dishes (P35G-1.5-14-C-GRID) were used to seed and infect HFF-1 cells with transiently transfected parasites that express pTUB-YFP-FAPP1PH-Myc or pTUB-YFP-SidM-P4M-Myc. At 24 hours post inoculation coverslips were fixed with 4% PFA and 0.025% glutaraldehyde (GA) in PBS for 15 min and washed 3X with PBS with a 5 min incubation per wash. Parasites were permeabilized with 0.2% Triton-X (v/v) in PBS for 4 min, washed 1X with PB (0.1 M phosphate buffer pH 7.2), incubated with primary mouse anti-GRA3 antibody secondary antibody directed against Alexa fluor 594 (A11032) (see **Supplementary Table 2** for used dilutions), stained with Hoechst solution (1:5000) for 5 min, and washed 3X with PB with a 10 min incubation per wash. Intracellular parasites were identified using a confocal NikonA1R microscope outfitted with a CFI Plan Apo lambda 60x/1.4 oil objective (to acquire z-sections of parasites with a LAP body to create a map of the parasite location), and a CFI Plan Apo lambda 20x/0.75 objective (to identify the position of parasites with a LAP body within the coordinate system in the glass-gridded coverslip).

Coordinates of interest in the grid were marked with nail varnish on the base of the coverslip. Cells were fixed again with 4% PFA and 0.5% GA in PB overnight at 4 &C, washed 3X with distilled water (10 min each), post-stained with 1% osmium (OsO_4_) for 15 min and with filtered 2% uranyl acetate for 20 min (water washes between each step – 3X and 10 min incubation each). The samples were dehydrated in a graded ethanol series (30%, 50%, 70%, 80%, 90%, 96%, 100%, 100%, 100%) with incubations of 10 min at each step. Cells on coverslips were embedded in a modified Quetol/Spurr’s resin mixed 1:1 with ethanol for 2 hours, followed by resin alone for 4 hours. To limit embedding to the targeted region of the glass-gridded coverslip, a plastic capsule (both ends open) was placed up-side-down on the coverslip over the varnish mark and incubated overnight at 60°C (Hanson et al., 2010). Immobilized capsules were filled with resin and incubated at 60°C for 48 hours. Resin blocks were detached by heating the coverslip with a passing flame for 7 sec under the bottom of the coverslip (Loussert et al., 2012). Serial ultrathin sections (∼100 nm) were collected using a Leica UC7 ultramicrotome and mounted on formvar slot grids. Sections were stained with lead citrate for 7-10 min and imaged using an FEI Morgagni 268 transmission electron microscope at 70 KeV with a MegaView III CCD camera and iTEM image acquisition software. Image analyses were performed with ImageJ. Data were collected from eight parasites per experimental condition.

### Statistical analyses

Data obtained from at least three independent biological replicates were presented as mean ± standard error of the mean (SEM). Two-way ANOVA followed by Dunnett’s multiple comparison test was used to compare two or more experimental groups to a single control group. One-way ANOVA followed by Tukey’s multiple comparison test was used to compare two or more experimental groups to every other group. T-test (Mann-Whitney test) was performed to compare the means of two groups. Data were calculated and illustrated using GraphPad Prism 6 software. A P value < 0.05 was set as threshold for statistical significance.

## ACKNOWLEDGMENTS

We thank Dr. Kenton Arkill (Univ. Nottingham, UK) for his expert advice on the optimization and imaging analysis of CLEM acquisitions. Sample preparation for EM imaging was performed at the Texas A&M University Microscopy and Imaging Center Core Facility (RRID:SCR_022128) with the help of Dr. Stanislav Vitha. Microscopy work performed using the Zeiss LSM 780 NLO multiphoton microscope provided with the Airyscan detector system and on the FEI Morgagni 268 transmission electron microscope at the Texas A&M School of Veterinary Medicine & Biomedical Sciences Image Analysis Laboratory with the assistance of Drs. Robert Burghardt and Joseph Szule. This work was supported by National Institutes of Health grant R35 GM131804 and BE0017 from the Robert A. Welch Foundation to VAB. The authors declare no competing interests.

## SUPPLEMENTARY MATERIAL

**Supplementary Table 1.**
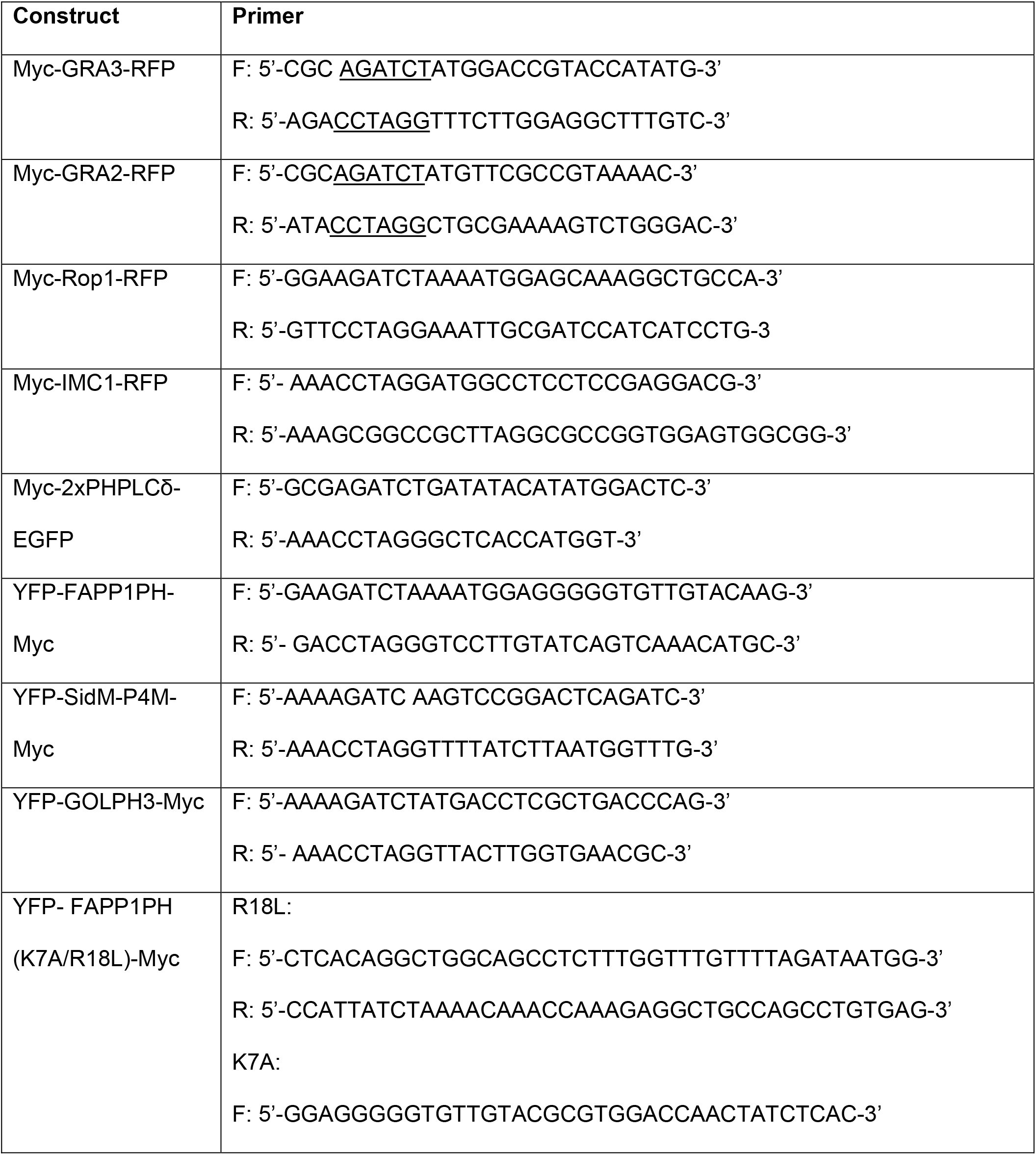
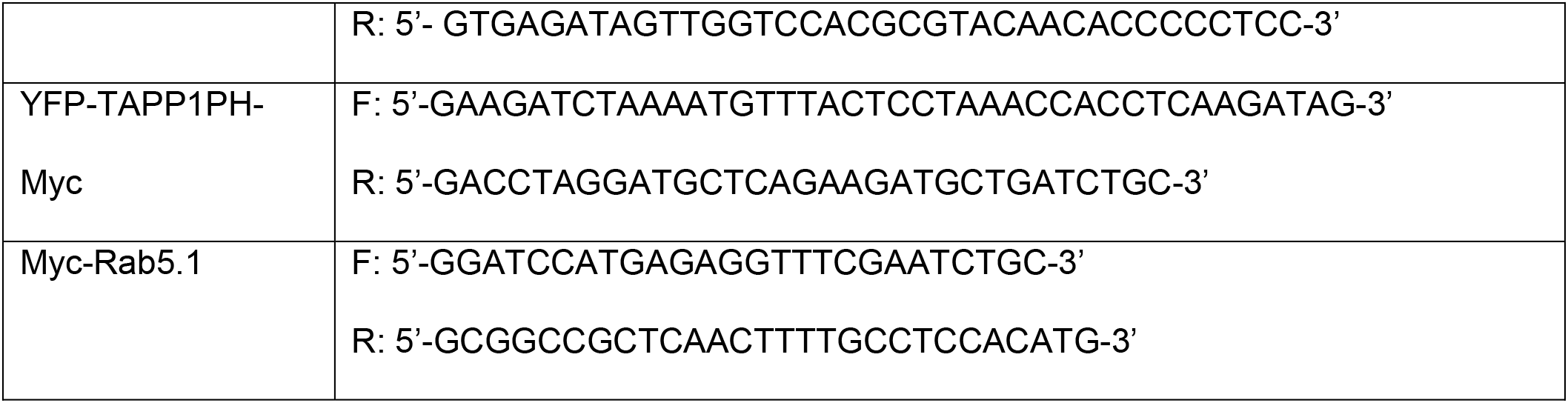
Primers used for subcloning *T. gondii* intracellular organelle marker genes used into the pTub expression vector.

**Supplementary Table 2.**
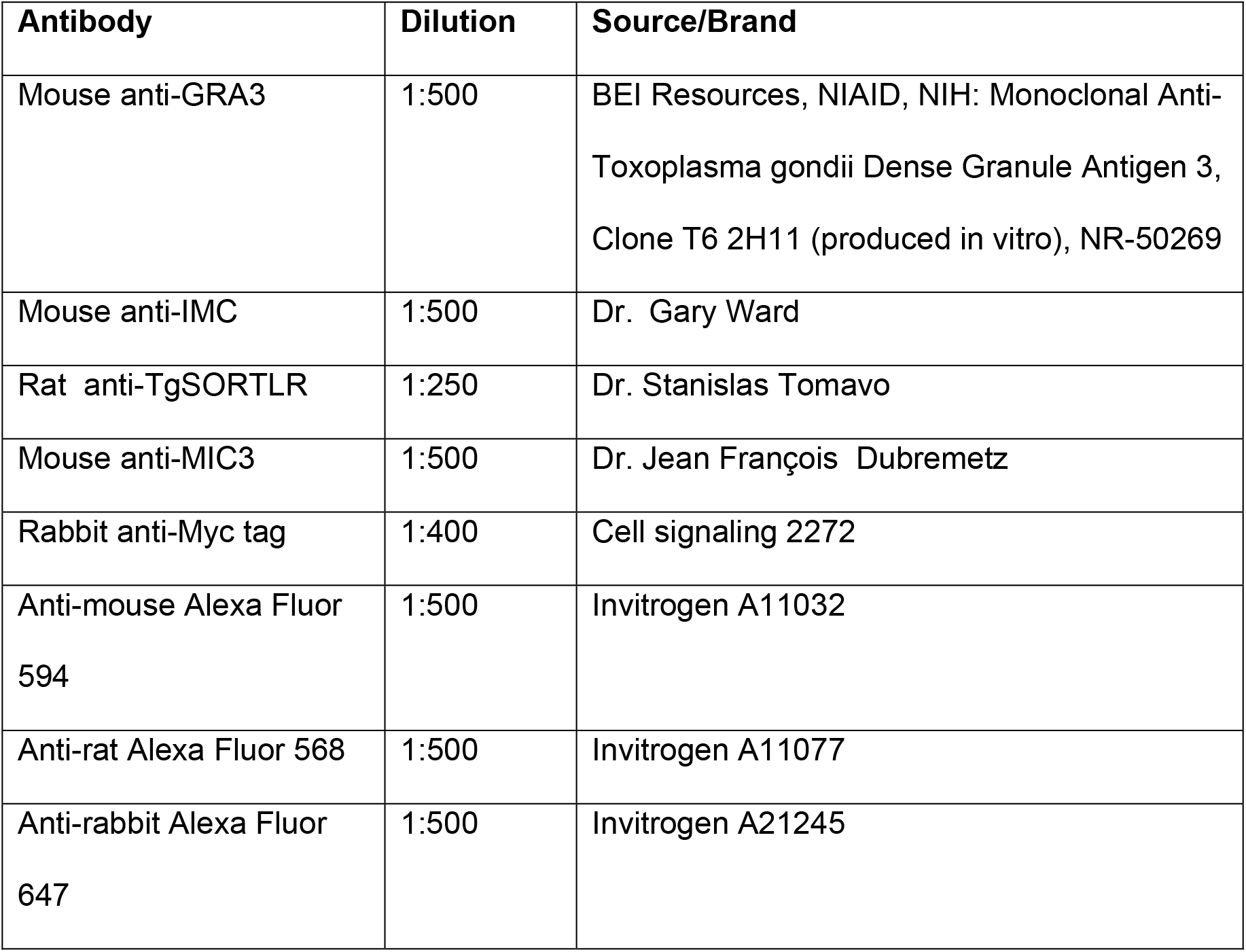
List of antibodies.

## REFERENCES

Bai, M.J., Wang, J.L., Elsheikha, H.M., Liang, Q.L., Chen, K., Nie, L.B. and Zhu, X.Q., 2018. Functional characterization of dense granule proteins in Toxoplasma gondii RH strain using CRISPR-Cas9 system. Frontiers in cellular and infection microbiology, 8, p.300.

Balla, T., 2013. Phosphoinositides: tiny lipids with giant impact on cell regulation. Physiological reviews, 93(3), pp.1019–1137.

Bankaitis, V.A., Aitken, J.R., Cleves, A.E. and Dowhan, W., 1990. An essential role for a phospholipid transfer protein in yeast Golgi function. Nature, 347(6293), pp.561–562.

Bankaitis, V.A., Mousley, C.J. and Schaaf, G., 2010. The Sec14 superfamily and mechanisms for crosstalk between lipid metabolism and lipid signaling. Trends in biochemical sciences, 35(3), pp.150–160.

Carpenter, C.L. and Cantley, L.C., 1990. Phosphoinositide kinases. Biochemistry, 29(51), pp.11147–11156.

Carruthers, V.B., Giddings, O.K. and Sibley, L.D., 1999. Secretion of micronemal proteins is associated with Toxoplasma invasion of host cells. Cellular microbiology, 1(3), pp.225–235.

Chaturvedi, S., Qi, H., Coleman, D., Rodriguez, A., Hanson, P.I., Striepen, B., Roos, D.S. and Joiner, K.A., 1999. Constitutive calcium-independent release of Toxoplasma gondii dense granules occurs through the NSF/SNAP/SNARE/Rab machinery. Journal of Biological Chemistry, 274(4), pp.2424–2431.

Clayton, E.L., Minogue, S. and Waugh, M.G., 2013. Phosphatidylinositol 4-kinases and PI4P metabolism in the nervous system: roles in psychiatric and neurological diseases. Molecular neurobiology, 47(1), pp.361–372.

Coppens, I., Andries, M., Liu, J.L. and Cesbron-Delauw, M.F., 1999. Intracellular trafficking of dense granule proteins in Toxoplasma gondii and experimental evidences for a regulated exocytosis. European journal of cell biology, 78(7), pp.463–472.

Daher, W., Morlon-Guyot, J., Sheiner, L., Lentini, G., Berry, L., Tawk, L., Dubremetz, J.F., Wengelnik, K., Striepen, B. and Lebrun, M., 2015. Lipid kinases are essential for apicoplast homeostasis in Toxoplasma gondii. Cellular microbiology, 17(4), pp.559–578.

Davidson, H.W., Rhodes, C.J. and Hutton, J.C., 1988. Intraorganellar calcium and pH control proinsulin cleavage in the pancreatic β cell via two distinct site-specific endopeptidases. Nature, 333(6168), pp.93–96.

Del Campo, C.M., Mishra, A.K., Wang, Y.H., Roy, C.R., Janmey, P.A. and Lambright, D.G., 2014. Structural basis for PI (4) P-specific membrane recruitment of the Legionella pneumophila effector DrrA/SidM. Structure, 22(3), pp.397–408.

Di Paolo, G. and De Camilli, P., 2006. Phosphoinositides in cell regulation and membrane dynamics. Nature, 443(7112), pp.651–657.

Dowler, S., Currie, R.A., Campbell, D.G., Deak, M., Kular, G., Downes, C.P. and Alessi, D.R., 2000. Identification of pleckstrin-homology-domain-containing proteins with novel phosphoinositide-binding specificities. Biochemical Journal, 351(1), pp.19–31.

Duex, J.E., Nau, J.J., Kauffman, E.J. and Weisman, L.S., 2006. Phosphoinositide 5-phosphatase Fig4p is required for both acute rise and subsequent fall in stress-induced phosphatidylinositol 3, 5-bisphosphate levels. Eukaryotic cell, 5(4), pp.723–731.

Dubey, J.P., Lindsay, D.S. and Speer, C., 1998. Structures of Toxoplasma gondii tachyzoites, bradyzoites, and sporozoites and biology and development of tissue cysts. Clinical microbiology reviews, 11(2), pp.267–299.

Escalante, A.A. and Ayala, F.J., 1995. Evolutionary origin of Plasmodium and other Apicomplexa based on rRNA genes. Proceedings of the National Academy of Sciences, 92(13), pp.5793–5797.

Gillooly, D.J., Morrow, I.C., Lindsay, M., Gould, R., Bryant, N.J., Gaullier, J.M., Parton, R.G. and Stenmark, H., 2000. Localization of phosphatidylinositol 3-phosphate in yeast and mammalian cells. The EMBO journal, 19(17), pp.4577–4588.

Grabon, A., Bankaitis, V.A. and McDermott, M.I., 2019. The interface between phosphatidylinositol transfer protein function and phosphoinositide signaling in higher eukaryotes. Journal of lipid research, 60(2), pp.242–268.

Griffith, M.B., Pearce, C.S. and Heaslip, A.T., 2022. Dense granule biogenesis, secretion, and function in Toxoplasma gondii. Journal of Eukaryotic Microbiology, p.e12904.

Halet, G., Tunwell, R., Balla, T., Swann, K. and Carroll, J., 2002. The dynamics of plasma membrane PtdIns (4, 5) P 2 at fertilization of mouse eggs. Journal of Cell Science, 115(10), pp.2139–2149.

Hama, H., Schnieders, E.A., Thorner, J., Takemoto, J.Y. and DeWald, D.B., 1999. Direct involvement of phosphatidylinositol 4-phosphate in secretion in the yeast Saccharomyces cerevisiae. Journal of Biological Chemistry, 274(48), pp.34294–34300.

Hammond, G.R., Fischer, M.J., Anderson, K.E., Holdich, J., Koteci, A., Balla, T. and Irvine, R.F., 2012. PI4P and PI (4, 5) P2 are essential but independent lipid determinants of membrane identity. Science, 337(6095), pp.727–730.

Hammond, G.R., Machner, M.P. and Balla, T., 2014. A novel probe for phosphatidylinositol 4-phosphate reveals multiple pools beyond the Golgi. Journal of Cell Biology, 205(1), pp.113–126.

Heo, W.D., Inoue, T., Park, W.S., Kim, M.L., Park, B.O., Wandless, T.J. and Meyer, T., 2006. PI (3, 4, 5) P3 and PI (4, 5) P2 lipids target proteins with polybasic clusters to the plasma membrane. Science, 314(5804), pp.1458–1461.

Hanson, H.H., Reilly, J.E., Lee, R., Janssen, W.G. and Phillips, G.R., 2010. Streamlined embedding of cell monolayers on gridded glass-bottom imaging dishes for correlative light and electron microscopy. Microscopy and Microanalysis, 16(6), pp.747–754.

He, J., Scott, J.L., Heroux, A., Roy, S., Lenoir, M., Overduin, M., Stahelin, R.V. and Kutateladze, T.G., 2011. Molecular basis of phosphatidylinositol 4-phosphate and ARF1 GTPase recognition by the FAPP1 pleckstrin homology (PH) domain. Journal of Biological Chemistry, 286(21), pp.18650–18657.

Heaslip, A.T., Nelson, S.R. and Warshaw, D.M., 2016. Dense granule trafficking in Toxoplasma gondii requires a unique class 27 myosin and actin filaments. Molecular biology of the cell, 27(13), pp.2080–2089.

Herneisen, A.L., Li, Z.H., Chan, A.W., Moreno, S.N. and Lourido, S., 2022. Temporal and thermal profiling of the Toxoplasma proteome implicates parasite Protein Phosphatase 1 in the regulation of Ca2+-responsive pathways. Elife, 11, p.e80336.

Hogan, A., Yakubchyk, Y., Chabot, J., Obagi, C., Daher, E., Maekawa, K. and Gee, S.H., 2004. The phosphoinositol 3, 4-bisphosphate-binding protein TAPP1 interacts with syntrophins and regulates actin cytoskeletal organization. Journal of Biological Chemistry, 279(51), pp.53717–53724.

Huff, J., 2015. The Airyscan detector from ZEISS: confocal imaging with improved signal-to-noise ratio and super-resolution. Nature methods, 12(12), pp.i–ii.

Ikonomov, O.C., Sbrissa, D. and Shisheva, A., 2006. Localized PtdIns 3, 5-P2 synthesis to regulate early endosome dynamics and fusion. American Journal of Physiology-Cell Physiology, 291(2), pp.C393–C404.

Jacot, D. and Soldati-Favre, D., 2020. CRISPR/Cas9-Mediated Generation of Tetracycline Repressor-Based Inducible Knockdown in Toxoplasma gondii. In Methods in Molecular Biology (pp. 125–141). Humana, New York, NY.

Jović, M., Kean, M.J., Szentpetery, Z., Polevoy, G., Gingras, A.C., Brill, J.A. and Balla, T., 2012. Two phosphatidylinositol 4-kinases control lysosomal delivery of the Gaucher disease enzyme, β-glucocerebrosidase. Molecular biology of the cell, 23(8), pp.1533–1545.

Jung, J.Y., Kim, Y.W., Kwak, J.M., Hwang, J.U., Young, J., Schroeder, J.I., Hwang, I. and Lee, Y., 2002. Phosphatidylinositol 3-and 4-phosphate are required for normal stomatal movements. The Plant Cell, 14(10), pp.2399–2412.

Karsten, V., Qi, H., Beckers, C.J., Reddy, A., Dubremetz, J.F., Webster, P. and Joiner, K.A., 1998. The protozoan parasite Toxoplasma gondii targets proteins to dense granules and the vacuolar space using both conserved and unusual mechanisms. The Journal of cell biology, 141(6), pp.1323–1333.

Kim, K., Soldati, D. and Boothroyd, J.C., 1993. Gene replacement in Toxoplasma gondii with chloramphenicol acetyltransferase as selectable marker. Science, 262(5135), pp.911–914.

Levine, N.D., 2018. The Protozoan Phylum Apicomplexa: Volume 2. CRC Press.

Levine, T.P. and Munro, S., 2002. Targeting of Golgi-specific pleckstrin homology domains involves both PtdIns 4-kinase-dependent and-independent components. Current Biology, 12(9), pp.695–704.

Li, W., Grech, J., Stortz, J.F., Gow, M., Periz, J., Meissner, M. and Jimenez-Ruiz, E., 2022. A splitCas9 phenotypic screen in Toxoplasma gondii identifies proteins involved in host cell egress and invasion. Nature Microbiology, 7(6), pp.882–895.

Ling, Y., Hayano, S. and Novick, P., 2014. Osh4p is needed to reduce the level of phosphatidylinositol-4-phosphate on secretory vesicles as they mature. Molecular biology of the cell, 25(21), pp.3389–3400.

Loussert, C., Forestier, C.L. and Humbel, B.M., 2012. Correlative light and electron microscopy in parasite research. In Methods in cell biology (Vol. 111, pp. 59–73). Academic Press.

Martins-Duarte, É.S., Sheiner, L., Reiff, S.B., de Souza, W. and Striepen, B., 2021. Replication and partitioning of the apicoplast genome of Toxoplasma gondii is linked to the cell cycle and requires DNA polymerase and gyrase. International journal for parasitology, 51(6), pp.493–504.

McNamara, C.W., Lee, M., Lim, C.S., Lim, S.H., Roland, J., Nagle, A., Simon, O., Yeung, B.K., Chatterjee, A.K., McCormack, S.L. and Manary, M.J., 2013. Targeting Plasmodium PI (4) K to eliminate malaria. Nature, 504(7479), pp.248–253.

Nguyen, P.M., Gandasi, N.R., Xie, B., Sugahara, S., Xu, Y. and Idevall-Hagren, O., 2019. The PI (4) P phosphatase Sac2 controls insulin granule docking and release. Journal of Cell Biology, 218(11), pp.3714–3729.

Nofal, S.D., Dominicus, C., Broncel, M., Katris, N.J., Flynn, H.R., Arrizabalaga, G., Botté, C.Y., Invergo, B.M. and Treeck, M., 2022. A positive feedback loop mediates crosstalk between calcium, cyclic nucleotide and lipid signalling in calcium-induced Toxoplasma gondii egress. PLoS Pathogens, 18(10), p.e1010901.

Merighi, A., 2018. Costorage of high molecular weight neurotransmitters in large dense core vesicles of mammalian neurons. Frontiers in cellular neuroscience, 12, p.272.

Mizushima, N., Levine, B., Cuervo, A.M. and Klionsky, D.J., 2008. Autophagy fights disease through cellular self-digestion. Nature, 451(7182), pp.1069–1075.

Nichols, B.A., Chiappino, M.L. and O’Connor, G.R., 1983. Secretion from the rhoptries of Toxoplasma gondii during host-cell invasion. Journal of ultrastructure research, 83(1), pp.85–98.

Olayioye, M.A., Noll, B. and Hausser, A., 2019. Spatiotemporal control of intracellular membrane trafficking by Rho GTPases. Cells, 8(12), p.1478.

Olsen, H.L., Høy, M., Zhang, W., Bertorello, A.M., Bokvist, K., Capito, K., Efanov, A.M., Meister, B., Thams, P., Yang, S.N. and Rorsman, P., 2003. Phosphatidylinositol 4-kinase serves as a metabolic sensor and regulates priming of secretory granules in pancreatic β cells. Proceedings of the National Academy of Sciences, 100(9), pp.5187–5192.

Omar-Hmeadi, M. and Idevall-Hagren, O., 2021. Insulin granule biogenesis and exocytosis. Cellular and Molecular Life Sciences, 78(5), pp.1957–1970.

Orii, M., Tsuji, T., Ogasawara, Y. and Fujimoto, T., 2021. Transmembrane phospholipid translocation mediated by Atg9 is involved in autophagosome formation. Journal of Cell Biology, 220(3).

Ouologuem, D.T. and Roos, D.S., 2014. Dynamics of the Toxoplasma gondii inner membrane complex. Journal of cell science, 127(15), pp.3320–3330.

Pelletier, L., Stern, C.A., Pypaert, M., Sheff, D., Ngô, H.M., Roper, N., He, C.Y., Hu, K., Toomre, D., Coppens, I. and Roos, D.S., 2002. Golgi biogenesis in Toxoplasma gondii. Nature, 418(6897), pp.548–552.

Petiot, A., Fauré, J., Stenmark, H. and Gruenberg, J., 2003. PI3P signaling regulates receptor sorting but not transport in the endosomal pathway. The Journal of cell biology, 162(6), pp.971–979.

Pieperhoff, M.S., Schmitt, M., Ferguson, D.J. and Meissner, M., 2013. The role of clathrin in post-Golgi trafficking in Toxoplasma gondii. PLoS One, 8(10), p.e77620.

Pittman, K.J., Aliota, M.T. and Knoll, L.J., 2014. Dual transcriptional profiling of mice and Toxoplasma gondii during acute and chronic infection. BMC genomics, 15(1), pp.1–19.

Rhodes, C.J., Lucas, C.A., Mutkoski, R.L., Orci, L. and Halban, P.A., 1987. Stimulation by ATP of proinsulin to insulin conversion in isolated rat pancreatic islet secretory granules. Association with the ATP-dependent proton pump. Journal of Biological Chemistry, 262(22), pp.10712–10717.

Rivas, M.P., Kearns, B.G., Xie, Z., Guo, S., Sekar, M.C., Hosaka, K., Kagiwada, S., York, J.D. and Bankaitis, V.A., 1999. Pleiotropic alterations in lipid metabolism in yeast sac1 mutants: relationship to “bypass Sec14p” and inositol auxotrophy. Molecular biology of the cell, 10(7), pp.2235–2250.

Rothman, J.E., 1994. Mechanisms of intracellular protein transport. Nature, 372(6501), pp.55–63.

Schaaf, G., Ortlund, E.A., Tyeryar, K.R., Mousley, C.J., Ile, K.E., Garrett, T.A., Ren, J., Woolls, M.J., Raetz, C.R., Redinbo, M.R. and Bankaitis, V.A., 2008. Functional anatomy of phospholipid binding and regulation of phosphoinositide homeostasis by proteins of the sec14 superfamily. Molecular cell, 29(2), pp.191–206.

Schekman, R. and Orci, L., 1996. Coat proteins and vesicle budding. Science, 271(5255), pp.1526–1533.

Schmiedeberg, L., Skene, P., Deaton, A. and Bird, A., 2009. A temporal threshold for formaldehyde crosslinking and fixation. PLoS One, 4(2), p.e4636.

Schoonderwoert, V.T.G., Holthuis, J.C., Tanaka, S., Tooze, S.A. and Martens, G.J., 2000. Inhibition of the vacuolar H+-ATPase perturbs the transport, sorting, processing and release of regulated secretory proteins. European journal of biochemistry, 267(17), pp.5646–5654.

Sloves, P.J., Delhaye, S., Mouveaux, T., Werkmeister, E., Slomianny, C., Hovasse, A., Alayi, T.D., Callebaut, I., Gaji, R.Y., Schaeffer-Reiss, C. and Van Dorsselear, A., 2012. Toxoplasma sortilin-like receptor regulates protein transport and is essential for apical secretory organelle biogenesis and host infection. Cell host & microbe, 11(5), pp.515–527.

Soldati, D. and Boothroyd, J.C., 1993. Transient transfection and expression in the obligate intracellular parasite Toxoplasma gondii. Science, 260(5106), pp.349–352.

Striepen, B., He, C.Y., Matrajt, M., Soldati, D. and Roos, D.S., 1998. Expression, selection, and organellar targeting of the green fluorescent protein in Toxoplasma gondii. Molecular and biochemical parasitology, 92(2), pp.325–338.

Tandon, A., Bannykh, S., Kowalchyk, J.A., Banerjee, A., Martin, T.F. and Balch, W.E., 1998. Differential regulation of exocytosis by calcium and CAPS in semi-intact synaptosomes. Neuron, 21(1), pp.147–154.

Tawk, L., Dubremetz, J.F., Montcourrier, P., Chicanne, G., Merezegue, F., Richard, V., Payrastre, B., Meissner, M., Vial, H.J., Roy, C. and Wengelnik, K., 2011. Phosphatidylinositol 3-monophosphate is involved in toxoplasma apicoplast biogenesis. PLoS pathogens, 7(2), p.e1001286.

Várnai, P. and Balla, T., 1998. Visualization of phosphoinositides that bind pleckstrin homology domains: calcium-and agonist-induced dynamic changes and relationship to myo-[3H] inositol-labeled phosphoinositide pools. The Journal of cell biology, 143(2), pp.501–510.

Walch-Solimena, C. and Novick, P., 1999. The yeast phosphatidylinositol-4-OH kinase pik1 regulates secretion at the Golgi. Nature cell biology, 1(8), pp.523–525.

Whitley, P., Hinz, S. and Doughty, J., 2009. Arabidopsis FAB1/PIKfyve proteins are essential for development of viable pollen. Plant physiology, 151(4), pp.1812–1822.

Wood, C.S., Schmitz, K.R., Bessman, N.J., Setty, T.G., Ferguson, K.M. and Burd, C.G., 2009. PtdIns4 P recognition by Vps74/GOLPH3 links PtdIns 4-kinase signaling to retrograde Golgi trafficking. Journal of Cell Biology, 187(7), pp.967–975.

Xie, Z., Hur, S.K., Zhao, L., Abrams, C.S. and Bankaitis, V.A., 2018. A Golgi lipid signaling pathway controls apical Golgi distribution and cell polarity during neurogenesis. Developmental cell, 44(6), pp.725–740.

Yeh, Y.T., Sona, C., Yan, X., Pathak, A., McDermott, M.I., Xie, Z., Liu, L., Arunagiri, A., Wang, Y., Cazenave-Gassiot, A. and Ghosh, A., 2022. Restoration of PITPNA in Type 2 diabetic human islets reverses pancreatic beta-cell dysfunction. bioRxiv. 2022.07.06.498991; doi: https://doi.org/10.1101/2022.07.06.498991

Yu, S., Satoh, A., Pypaert, M., Mullen, K., Hay, J.C. and Ferro-Novick, S., 2006. mBet3p is required for homotypic COPII vesicle tethering in mammalian cells. The Journal of cell biology, 174(3), pp.359–368.

Zhang, X., Jiang, S., Mitok, K.A., Li, L., Attie, A.D. and Martin, T.F., 2017. BAIAP3, a C2 domain–containing Munc13 protein, controls the fate of dense-core vesicles in neuroendocrine cells. Journal of Cell Biology, 216(7), pp.2151–2166.

